# Molecular determinants of fibrillation in a viral amyloidogenic domain from combined biochemical and biophysical studies

**DOI:** 10.1101/2022.11.02.514851

**Authors:** Juliet F. Nilsson, Hakima Baroudi, Frank Gondelaud, Giulia Pesce, Christophe Bignon, Denis Ptchelkine, Joseph Chamieh, Hervé Cottet, Andrey V. Kajava, Sonia Longhi

## Abstract

The Nipah and Hendra viruses (NiV and HeV) are biosafety level 4 human pathogens classified within the *Henipavirus* genus of the *Paramyxoviridae* family. In both NiV and HeV, the gene encoding the Phosphoprotein (P protein), an essential polymerase cofactor, also encodes the V and W proteins. These three proteins, which share an intrinsically disordered N-terminal domain (NTD) and have unique C-terminal domains (CTD), are all known to counteract the host innate immune response, with V and W acting by either counteracting or inhibiting Interferon (IFN) signaling. Recently, using a combination of biophysical and structural approaches, the ability of a short region within the shared NTD (PNT3 region) to form amyloid-like structures was reported. Here, we evaluated the relevance of each of three contiguous tyrosine residues located in a previously identified amyloidogenic motif (EYYY) within HeV PNT3 to the fibrillation process. Our results indicate that removal of a single tyrosine in this motif significantly decreases the ability to form fibrils independently of position, mainly affecting the elongation phase. In addition, we show that the C-terminal half of PNT3 has an inhibitory effect on fibril formation that may act as a molecular shield and could thus be a key domain in the regulation of PNT3 fibrillation. Finally, the kinetics of fibril formation for the two PNT3 variants with highest and lowest fibrillation propensity were studied by Taylor Dispersion Analysis (TDA). The results herein presented shed light onto the molecular mechanisms involved in fibril formation. In addition, the PNT3 variants we generated represent valuable tools to further explore the functional impact of V/W fibrillation in transfected and infected cells.

## 1. Introduction

Hendra virus (HeV), together with the closely related Nipah virus (NiV), is a Biosafety Level 4 (BSL-4) pathogen belonging to the *Henipavirus* genus within the *Paramyxoviridae* family [1]. Henipaviruses are zoonotic viruses responsible in humans for severe encephalitis [1]. They are enveloped viruses with a non-segmented, single-stranded RNA genome of negative polarity [2]. Their genome is wrapped by the nucleoprotein (N) within a helical nucleocapsid that is the template used by the viral polymerase for transcription and replication. The polymerase consists of the L protein, which bears all the enzymatic activities, and of the phosphoprotein (P). P serves as an indispensable polymerase co-factor as not only it tethers the L protein onto the nucleocapsid, but also keeps L in a soluble and competent form for transcription and replication [3-5].

The N and P proteins from henipaviruses encompass long intrinsically disordered regions (IDRs) [6-8], *i*.*e*. regions devoid of stable secondary and tertiary structure [9-12]. The *Henipavirus* P protein consists of a long N-terminal intrinsically disordered domain (NTD) and a C-terminal region that possesses both structured and disordered regions (**Fig. 1**) [7,8,13-18].

**Figure 1.**
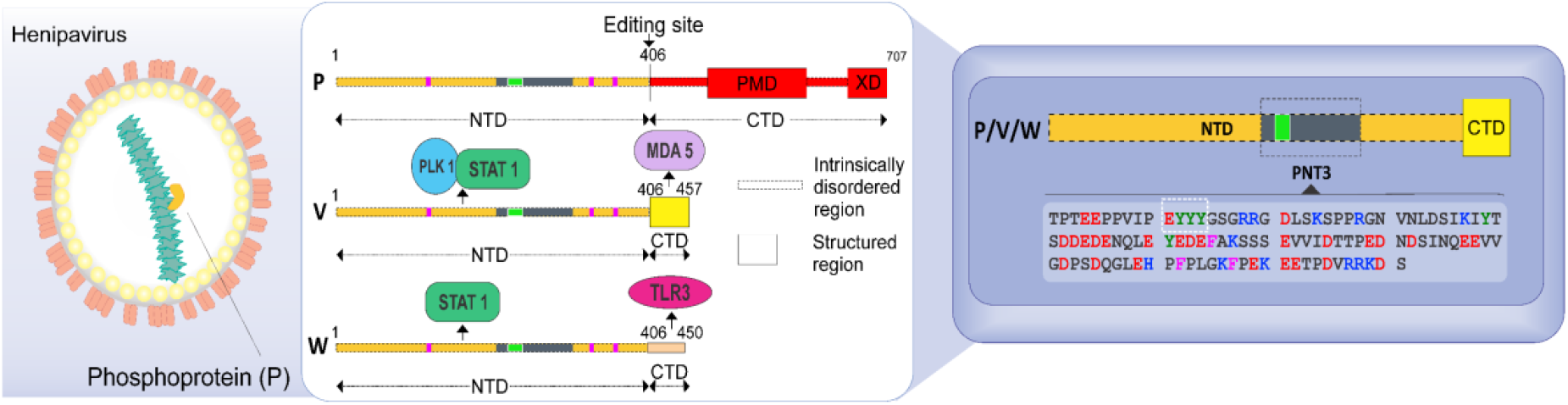
Schematic illustration of the HeV particle, of the organization of the P, V and W proteins and sequence of the HeV PNT3 region shared by the P, V, and W proteins. The left panel displays a scheme of the HeV virion, with the genome encapsidated by the nucleoprotein (in green) and the P (yellow) and the Large (beige) proteins attached onto the nucleocapsid. The central panel displays a scheme of the P, V and W protein organization, showing that they share a common N-terminal domain (NTD) and have distinct C-terminal domains (CTD). PMD: P multimerization domain; XD: X domain of P. Interaction sites with proteins associated with the host innate immune response are shown. STAT: Transducers and Activators of Transcription; PLK1: Polo-Like Kinase 1; MDA5: Melanoma Differentiation-Associated protein 5; TLR3: Toll-like receptor 3. Pink bars correspond to cysteine residues within NTD. The localization and the sequence of the PNT3 region within the NTD is shown in grey and its sequence is displayed in the blue, right panel. The EYYY motif is shown as a green square and is underlined within the PNT3 sequence.

As in many paramyxoviruses [19], the P gene from HeV and NiV also encodes the C, V and W non-structural proteins. While the C protein is encoded from an alternative reading frame within the P gene, the V and W proteins (∼50 kDa) result from a mechanism of co-transcriptional editing of the P messenger: the addition of either one or two non-templated guanosines at the editing site of the P messenger yields the V and W proteins, respectively (**Fig. 1**). The editing site is located at the end of the NTD-encoding region (**Fig, 1**). The P, V and W proteins therefore share a common NTD but have distinct C-terminal domains (CTDs) (**Fig. 1**). While the CTD of V adopts a zinc-finger conformation [20], the CTD of W is disordered [21].

The V and W proteins are key players in the evasion of the antiviral type I interferon (IFN-I)-mediated response [22-24]. This property relies on their ability to bind to a number of key cellular proteins involved in the antiviral response (**Fig. 1**).

We previously reported the ability of the HeV V protein to undergo a liquid-to-gel transition, with a region within the NTD (referred to as PNT3, aa 200-310) (**Fig. 1**) being identified as responsible for this behavior [25]. In those previous studies, we characterized PNT3 using a combination of biophysical and structural approaches. Congo red binding assays, together with negative-staining transmission electron microscopy (ns-TEM) studies, showed that PNT3 forms amyloid-like structures [25]. Noteworthy, Congo red staining experiments provided hints that these amyloid-like fibrils form not only *in vitro* but also *in cellula* after transfection or infection suggesting a probable functional role. In light of the critical role of the *Henipavirus* V and W proteins in evading the host innate immune response, we previously proposed that in infected cells PNT3-mediated fibrillar aggregates could sequester key cellular proteins involved in the antiviral response such as such as STAT and 14-3-3 proteins, thereby preventing IFN signaling and abrogating NF-κB-induced proinflammatory response [26]. Consistent with the presence of PNT3, the *Henipavirus* W proteins were shown to be able to form amyloid-like fibrils as well [21].

Within PNT3, a motif encompassing three contiguous tyrosines (EYYY) was predicted as an amyloidogenic region [25]. The ArchCandy predictor [27,28] also predicted a fibril architecture in which the three contiguous tyrosines of the motif are part of the first β-strand of a β-strand-loop-β-strand motif (see Fig. 4C in [25]). ArchCandy identifies amyloidogenic regions based on their ability to form β-arcades. Indeed, the core structural element of a majority of naturally-occurring and disease-related amyloid fibrils is a β-arcade representing a parallel and in register stacks of β-strand-loop-β-strand motifs called β-arches [27]. Substitution of the three contiguous tyrosine residues with three alanine residues yielded a variant (referred to as PNT3^3A^) that was shown to possess a dramatically reduced fibrillation ability, thus providing direct experimental evidence for the predicted involvement of the EYYY motif in building up the core of the fibrils [25].

Here, with the aim of achieving a better understanding of the molecular determinants of HeV PNT3 fibrillation, we designed and characterized a set of additional PNT3 variants that were conceived to either further investigate the EYYY amyloidogenic motif or probe the contribution of the C-terminal half of the protein to the fibrillation process. Results, as obtained by combining various biophysical and structural approaches, show that removal of one out of the three tyrosines of the motif, irrespective of position, is sufficient to lead to a significantly reduced fibrillation ability. In addition, our results revealed that the C-terminal half of PNT3 acts as a natural dampener of the fibrillation process.

## 2. Results and Discussion

### 2.1. Influence of pH on the formation of HeV PNT3 amyloid-like fibrils

We previously documented the ability of the PNT3 region of the HeV V protein to form amyloid-like structures [25]. A number of studies reported an impact of pH on both fibril structure and kinetics [29-31]. As a first step towards an in-depth characterization of PNT3 fibrils, and with the aim of selecting appropriate conditions to investigate the kinetics of fibril formation, we sought at assessing the possible impact of pH on the fibrillation process. To this end, we used a previously described method based on the titration of polyethylene glycol (PEG), a crowding agent, to quantitatively assess the relative solubility of proteins [32]. Hence, after optimization of this method (see Materials and Methods), we performed PEG precipitation assays to evaluate the relative solubility of HeV PNT3 at three different pH values, namely 6.5, 7.2 and 8.0 (**Fig. 2**). From this assay, the PEG_1/2_ value, which corresponds to the PEG concentration at which 50% of the protein is still soluble, can be obtained and allows comparing protein aggregation propensities under different conditions [32]. Results display that HeV PNT3 at pH 6.5 shows less relative solubility compared to pH 7.2, indicating a higher aggregation propensity at the lower pH (**Fig. 2A**). This behavior might be at least partly rationalized based on the isoelectric point of the protein (pI=4.6), as proteins are well known to display minimal solubility at pH values close to their pI. The impact of pH on fibril formation, and the correlation with the PEG_1/2_ value, was confirmed by ns-TEM (**Fig. 2B**). The obtained micrographs show that at pH 6.5 fibrillar aggregates are present even at time 0, while at pH 7.2 equivalent fibrillar aggregates are only observed after an incubation of 96 hours at 37 °C (*i.e*. no fibrillar aggregates can be detected at time 0, **Fig. 2B**). This trend is further confirmed at pH 8, a condition where PNT3 displays the lowest propensity to form fibrillar aggregates (**Fig. 2**). Notably, in addition to the presence of fibrils, the micrographs obtained at the three pH values also show the presence of amorphous aggregates. These results, beyond advocating for a role of electrostatics in the aggregation process, prompted us to define pH 7.2 as the standard pH value for further studies: the rationale for this choice was that we wanted to be able to monitor the appearance and growth of fibrils over time, while at pH 6.5 fibrils were detected as early as at time zero.

**Figure 2.**
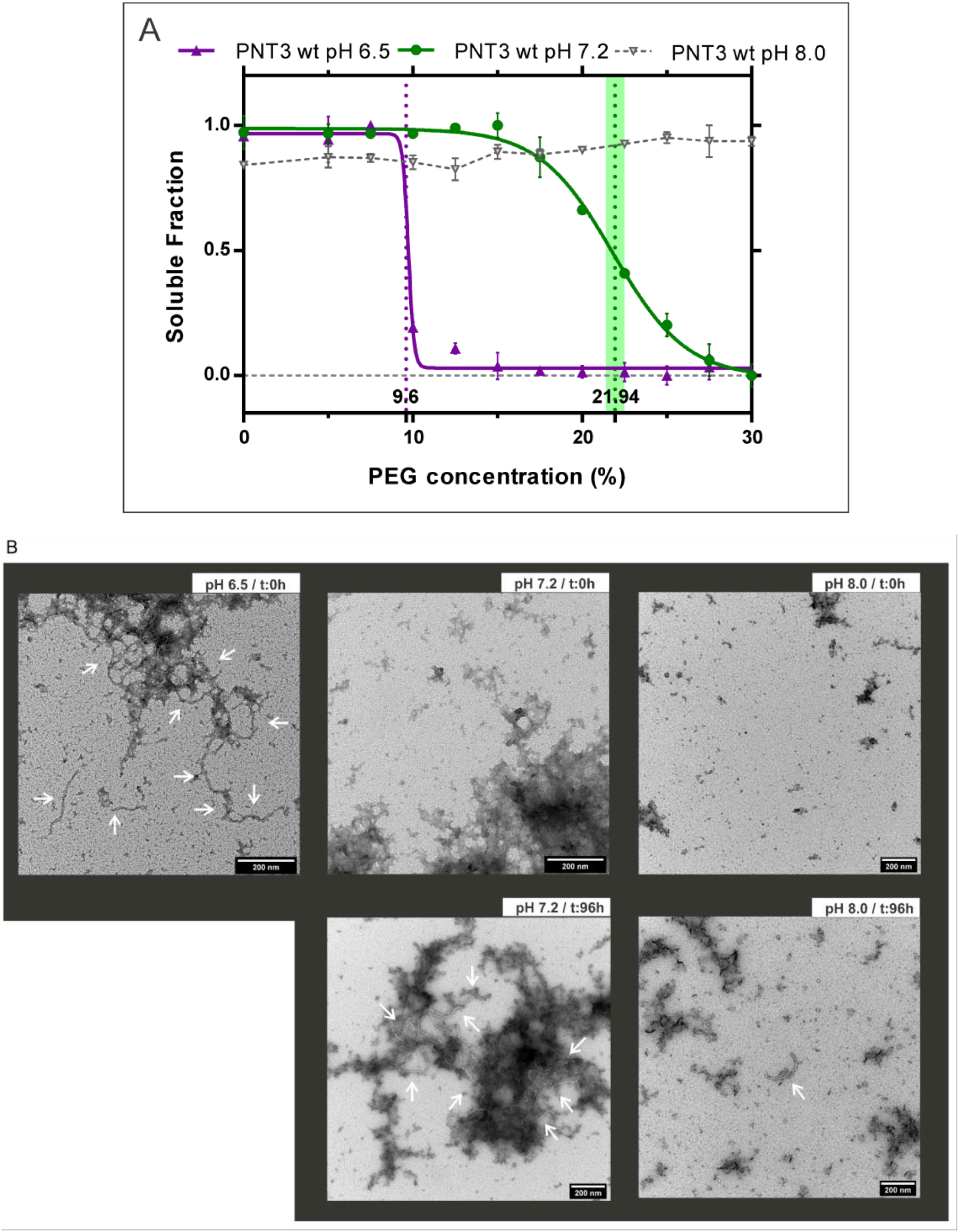
Fibrillation propensity of HeV PNT3 *wild-type* (*wt*) at different pHs. **A**. PEG assay and relative solubility of HeV PNT3 *wt* at pH 7.2 (green), 6.5 (violet) and 8.0 (light gray). The vertical lines correspond to PEG_1/2_ values (with their 95 % confidence intervals) as obtained after a normalization and fitting step to a sigmoid function. Note that the curve obtained at pH 6.5 exhibits poor fitting thus preventing calculation of the confidence interval. Data points at pH 8.0 could obviously not be fitted. **B**. Ns-TEM of HeV PNT3 *wt* fibrils at three different pH values: 6.5 (at time 0, t:0h), 7.2 and 8.0 (at time 0 and after 96 hours of incubation at 37 °C, t:96h). White arrows indicate fibrils.

### 2.2. Rational design and generation of PNT3 variants

#### 2.2.1. Design of PNT3 variants targeting the EYYY motif

Within the HeV PNT3 region, we previously identified an amyloidogenic motif encompassing three contiguous tyrosine residues (EYYY) [25]. The relevance of this motif for fibril formation was experimentally confirmed through the generation of the PNT3^3A^ variant, in which the three tyrosine residues were replaced with three alanine residues, which showed a reduced ability to form amyloid-like fibrils [25]. With the goal of further investigating this amyloidogenic motif and of unveiling whether all the three tyrosine residues were critical for fibril formation or whether only a subset of them was so, we rationally designed three single-site PNT3 variants where each one of the three contiguous tyrosine residues was replaced with one alanine (PNT3^A1^, PNT3 ^A2^ and PNT3^A3^, **Fig. 3A**). In this context, a construct encoding the corresponding PNT3 region from the NiV V protein (NiV PNT3) was also generated taking advantage of the fact that the NiV PNT3 motif encompasses only 2 contiguous tyrosine residues (EHYY) (**Fig. 3A** and **Supplementary Fig. S1**).

**Figure 3.**
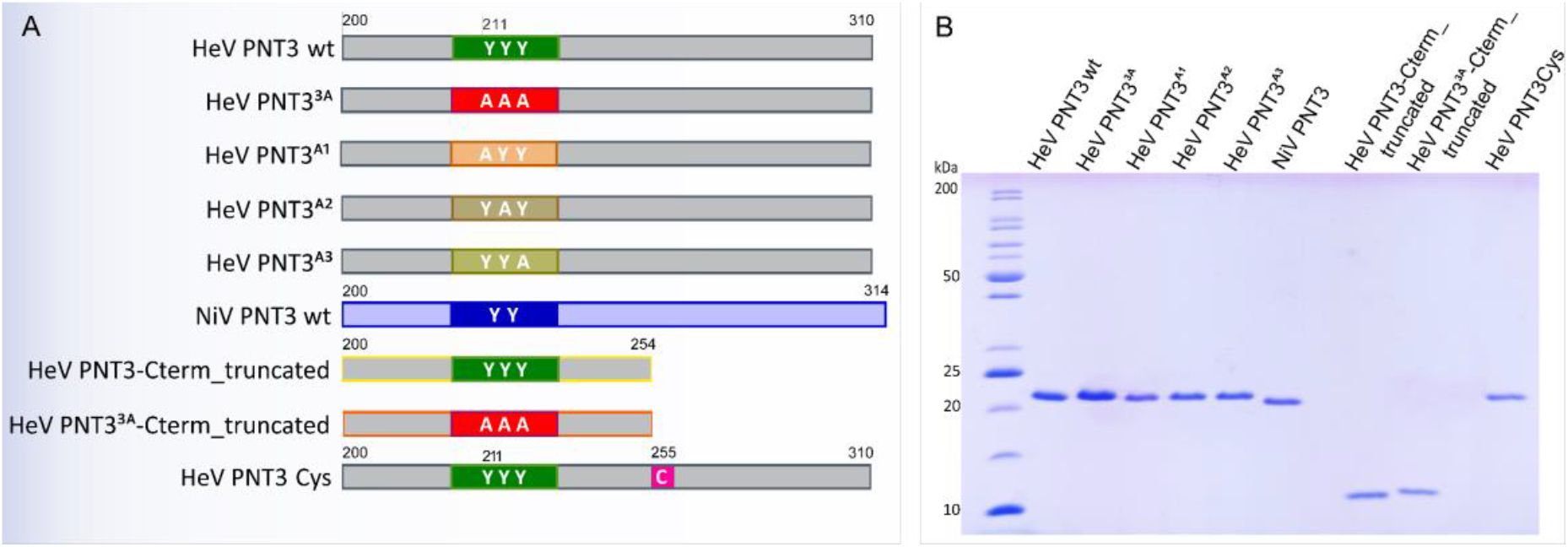
**A**. Schematic diagram of HeV PNT3 variants designed and constructed in this work. The EYYY motif of the HeV PNT3 is shows in green. **B**. SDS-PAGE analysis of purified proteins

#### 2.2.2. Design of HeV PNT3 truncated variants devoid of the C-terminal region

As mentioned above, the EYYY motif plays a significant role in fibril formation. However, the fact that the substitution of the triple tyrosine motif only reduces but does not fully abrogate the ability of PNT3 to form amyloid-like fibrils [25] indicates that other motifs and/or sequence attributes, that remained to be identified, contribute to the fibrillation process. Hence, to deepen the characterization and to assess the possible contribution of the PNT3 C-terminal region, we designed a C-terminally truncated HeV PNT3 variant (PNT3 C-term truncated) that lacks the second half of the protein (**Fig. 3A**). In addition, we also designed a C-terminally truncated HeV PNT3 variant where the three contiguous tyrosines of the EYYY motif were replaced with three alanines (PNT3^3A^ C-term truncated) (**Fig. 3A**).

#### 2.2.3. Design of a HeV PNT3 variant bearing a unique cysteine

The N-terminal domain, shared by the HeV P, V, and W proteins, has 3 cysteines distributed along the sequence (**Fig. 1**). Recently, Pesce & Gondelaud et. al suggested that disulfide bridges could be involved in preventing aggregation of the W protein [21]. Thus, in this context, we reasoned that a HeV PNT3 variant bearing a cysteine residue could be useful to investigate the possible impact of disulfide bridge-mediated protein dimerization on the fibrillation abilities of PNT3. We targeted for cysteine substitution the unique alanine residue of PNT3 (Ala255) to yield PNT3 variant A255C, PNT3_Cys (**Fig. 3A**). The rationale for choosing an alanine, rather than a serine residue which would have enabled a more isosteric substitution, was to introduce an as much conservative as possible substitution, while preserving the content in OH groups, which might play a role in the establishment of stabilizing inter-chain interactions in the core of the fibrils.

#### 2.2.4. Expression and purification of the PNT3 variants

All the proteins were expressed in *E. coli* as hexahistidine tagged forms with no solubility tag. The proteins were purified from the total fraction of the bacterial lysate under denaturing conditions. The proteins were purified by Immobilized Metal Affinity Chromatography (IMAC) and size exclusion chromatography (SEC). The purity of the final purified products was assessed by SDS-PAGE (**Fig. 3B**). The identity of all the variants was confirmed by mass spectrometry (MS) analysis of peptides resulting from the tryptic digestion of each protein (data not shown).

#### 2.2.5. Conformational characterization of the PNT3 variants

First, we evaluated the hydrodynamic properties of all the variants through analytical SEC. **Table 1** shows the Stokes radius (R_s_) value inferred for each protein, along with the Compaction Index (CI) associated to each variant. By comparing the mean measured Stokes radius (R_S_^OBS^) with the theoretical Stokes radii expected for various conformational states (*i.e*., Rs^NF^: natively folded protein; R_S_^PMG^: expected for a PMG; R_S_^U^: fully unfolded form; R_S_^IDP^: expected for an IDP), all the proteins were found to have R_S_ values consistent with a PMG state [33]. Their compaction indexes are relatively close to each other’s, with the notable exception of the HeV PNT3_C-term_truncated variant that is much more compact. Strikingly, the introduction of the triple alanine motif in the context of the truncated variant leads to a more extended conformation, a phenomenon already observed, although with a borderline significance, in the context of the full-length PNT3 protein (*cf*. HeV PNT3 *wt* and HeV PNT3^3A^ in **Table 1**). These results suggest that the second half of the protein is a determinant of chain expansion, with this effect being counteracted by the presence of the triple alanine motif.

**Table 1.**
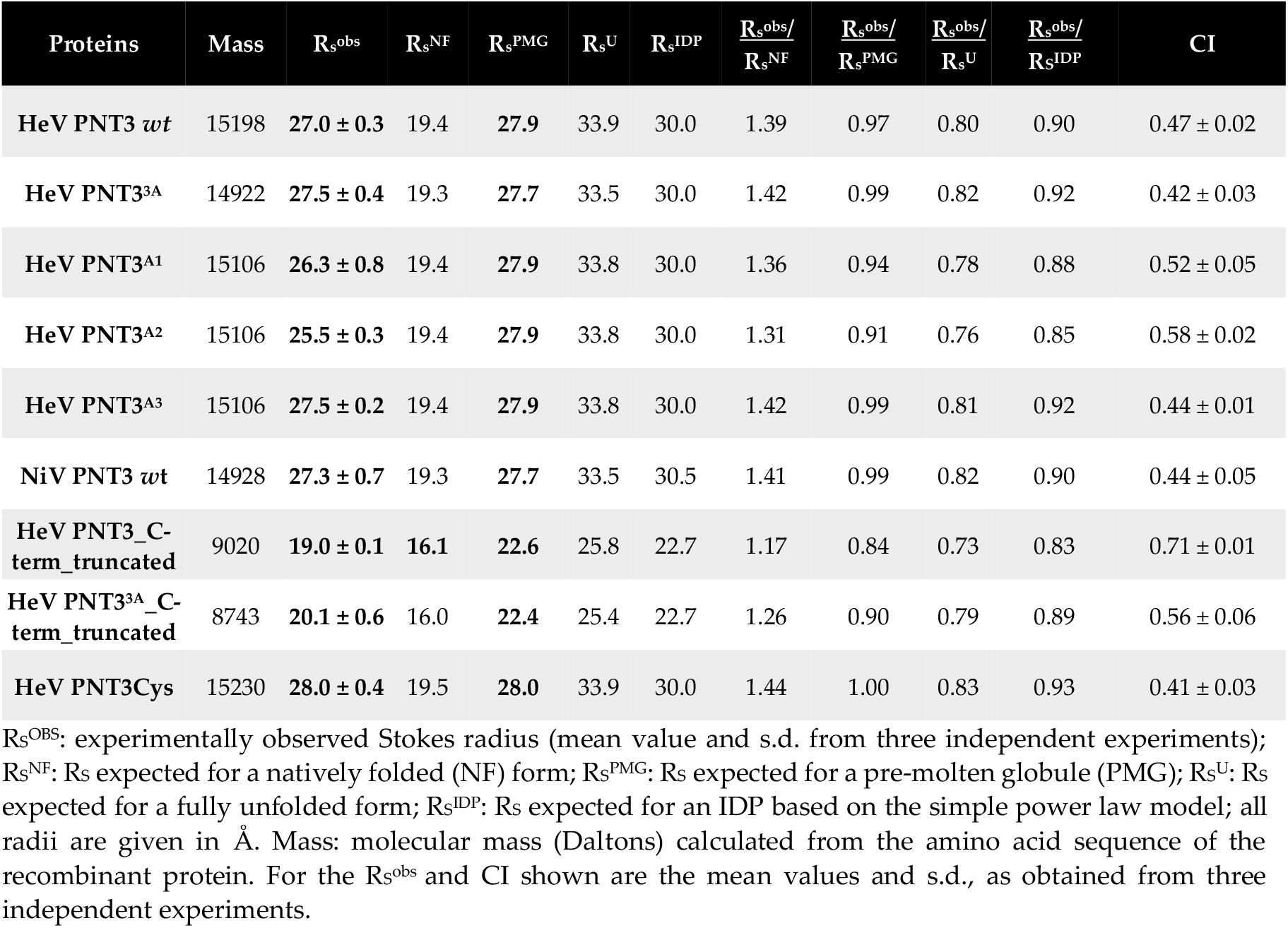
Stokes radii (R_S_^obs^, Å) of the PNT3 variants as inferred from the elution volume of the major SEC peak. Shown are also the expected values for the various conformational states, along with the ratios between the R^obs^ and each R_s_ state, and compaction index (CI) values.

In order to evaluate the secondary structure content of each variant we performed a Circular Dichroism (CD) analysis in the far ultraviolet (UV) region. All the variants present a spectrum typical of a disordered protein lacking any stable organized secondary structure, as judged from the large negative peak centered at 200 nm, and from the low ellipticity in the 220–230 nm region and at 190 nm (see [34] and references therein cited) (**Supplementary Fig. S2**). These results indicate that the introduced substitutions and/or the truncation impact only marginally, if at all, the secondary structure content of the protein. They also indicate that NiV PNT3 has a secondary structure content very close to that of its HeV counterpart. The finding that the CD spectra of PNT3 variants bearing the triple alanine motif are virtually superimposable onto those of variants bearing either the naturally occurring triple tyrosine motif or just one Tyr to Ala substitution, rules out the possibility that the expansion effect driven by the triple alanine motif, as observed in SEC studies, may arise from the presence of a transiently populated α-helix encompassing the motif. Therefore, the mechanism underlying the counteracting effect exerted by the triple alanine motif on chain compaction remains to be elucidated.

To achieve a more quantitative description of the conformational properties of the variants, we carried out Small-Angle X-ray Scattering (SAXS) studies coupled to SEC (SEC-SAXS). We selected a set of representative variants (*i.e*., HeV PNT3^3A^, NiV PNT3, HeV PNT3_C-term_truncated, and HeV PNT3^3A^_C-term_truncated) along with HeV PNT3 *wt*. Although we already previously reported SEC-SAXS studies of HeV PNT3 *wt* [25], this sample was herein again investigated under exactly the same conditions used for the other variants, so as to enable meaningful comparisons. For all the five PNT3 proteins, linearity of the Guinier region in the resulting scattering curves (**Supplementary Fig. S3A**) allowed meaningful estimations of the R_g_ (**Table 2**). The R_g_ values obtained for HeV PNT3^3A^, NiV PNT3, and HeV PNT3 *wt* are very close to each other’s clustering in a group with R_g_ around 37-39 Å. As expected, the two truncated variants have smaller, and close to each other’s, R_g_ values (**Table 2**). Notably, all experimental R_g_ obtained are close to the theoretical R_g_^U^, corresponding to chemically denatured (U) proteins, reflecting a highly extended conformation. Because of this, the R_g_-based CI could not be computed, the numerator (R_G_^U^ - R_G_^OBS^) in Eq. 9 being ≤ 0 (see Materials & Methods). These results are in contrast with the previous SEC results, where the variants were found to adopt a PMG conformation. A possible explanation for this might be related to the differences in the buffer used in the two techniques. As expected, the five variants were all found to be disordered as judged from the presence of a plateau in the normalized Kratky (**Supplementary Fig. S3B**) and the Kratky-Debye plots (**Supplementary Fig. S3C**). However, the two truncated variants, and particularly the HeV PNT3 C-term truncated one, showed a slight deviation from the pure random-coil regime as observed in the normalized Kratky plot. This deviation is consistent with a slightly more compact conformation, in line with the R_S_-based CI values discussed above.

**Table 2.** *R*_*g*_ and *D*_*max*_ as obtained from SEC-SAXS studies and expected values for the various conformational states.

| Proteins | $I(0) \text{ cm}^{-1}$ | $R_g \text{ (Å)}$<br>(Guinier) | $D_{max} \text{ (Å)}$ | $R_g^{IDP} \text{ (Å)}$ | $R_g^U \text{ (Å)}$ |
| --- | --- | --- | --- | --- | --- |
| HeV PNT3 *wt* | $0.03 \pm 1.8 \times 10^{-4}$ | $36.67 \pm 0.41$ | 140 | 32.6 | 35.9 |
| HeV PNT3<sup>3A</sup> | $0.022 \pm 8.1 \times 10^{-5}$ | $39.48 \pm 0.33$ | 144 | 32.6 | 35.9 |
| NiV PNT3 | $0.03 \pm 8.2 \times 10^{-5}$ | $37.37 \pm 0.21$ | 147 | 33.1 | 36.5 |
| HeV PNT3 C-term_truncated | $0.018 \pm 4.1 \times 10^{-5}$ | $27.53 \pm 0.14$ | 115 | 24.5 | 25.9 |
| HeV PNT3<sup>3A</sup> C-term_truncated | $0.015 \pm 5.6 \times 10^{-5}$ | $27.34 \pm 0.21$ | 119 | 24.5 | 25.9 |

$I(0)$ : Intensity at zero angle as determined from Guinier approximation; $R_g$ Guinier: $R_g$ values as obtained from Guinier approximation; $D_{max}$ : maximal intramolecular distance from $P(r)$ . $R_g^{IDP}$ : $R_g$ expected for an IDP based on the simple power-law model. $R_g^U$ : theoretical $R_g$ value expected for chemically denatured (U) proteins.

### 2.3. Relevance of the PNT3 EYYY motif in fibrillation abilities

#### 2.3.1. Aggregation propensity of the EYYY motif PNT3 variants

With the aim of elucidating the contribution of each tyrosine in the EYYY motif to the fibrillation process, we first assessed the aggregation propensity in the set of variants bearing alanine substitutions within the EYYY amyloidogenic motif. To this end, we took advantage of the same PEG solubility assay described above [32]. For each of the five EYYY variants (PNT3^3A^, PNT3^A1, A2, A3^ and NiV PNT3), we therefore carried out PEG solubility assays which enabled ranking them based on their estimated PEG_1/2_ value.

The PNT3^3A^ variant shows a significant increase in the relative solubility compared to PNT3 HeV *wt* (**Fig. 4A** and **4B**), a result in agreement with previous findings that pointed out a much lower propensity to form amyloid-like fibrils for PNT3^3A^ [25]. The three variants where only one Tyr was replaced, however, show no significant difference in the PEG_1/2_ values compared to HeV PNT3 *wt* (**Fig. 4B**), suggesting that the removal of one tyrosine does not affect the aggregation propensity of PNT3. By contrast, and interestingly, NiV PNT3 (EHYY) displays an intermediate relative solubility between HeV PNT3 *wt* and HeV PNT3^3A^ (**Fig. 4A**). In light of the results obtained with the HeV PNT3 variants bearing two tyrosines (*i.e.*, PNT3^A1^, PNT3^A2^ and PNT3^A3^), the intermediate aggregation propensity of NiV PNT3 more likely arise from differences in the amino acid context rather than from the absence of just one tyrosine.

**Figure 4.**
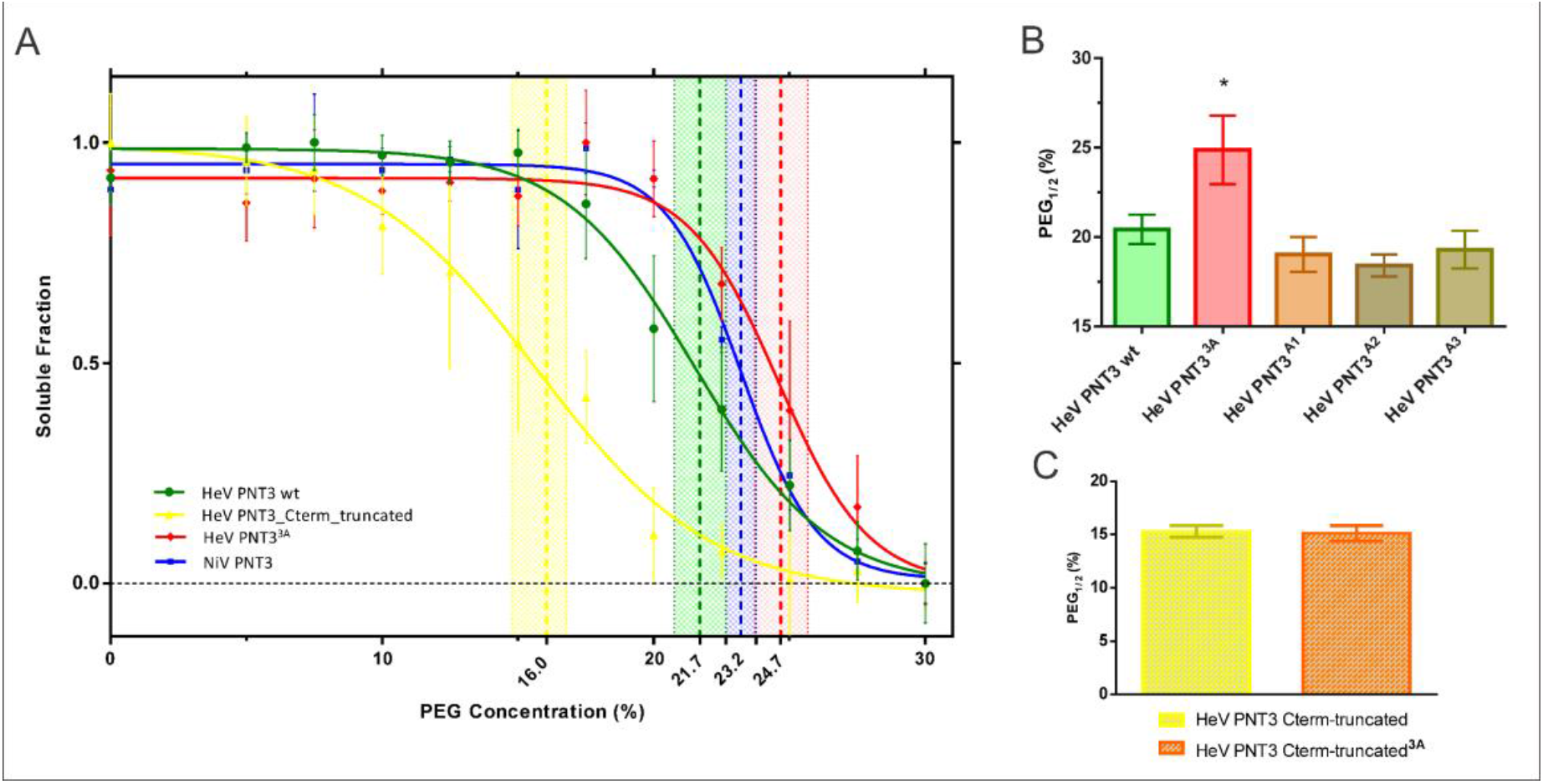
PEG solubility assay for the set of PNT3 variants. **A**. Soluble fraction of each variant at different PEG concentrations. HeV PNT3 *wt* (green), HeV PNT3^3A^ (red), HeV PNT3_C-term_truncated (yellow), NiV PNT3 (blue). Vertical lines represent PEG_1/2_ values including their 95 % confidence intervals obtained after a normalization and fitting step to a sigmoid function. **B**. PEG_1/2_ values obtained for the *wt*, the triple alanine variant and the single alanine HeV PNT3 variants. **C**. PEG_1/2_ values obtained for the C-terminally truncated variants.

#### 2.3.2. Congo Red binding abilities of PNT3 EYYY motif variants

Congo Red (CR) is a widely used dye to document the presence of amyloids: binding of this dye to cross β-sheet structures in fact leads to hyperchromicity and a red shift in the absorbance maximum of the CR spectrum. Hence, to further characterize PNT3 EYYY motif variants, we took advantage of CR binding assays. We compared the binding abilities of the PNT3^A1^, PNT3 ^A2^, PNT3^A3^ and NiV variants to those of both PNT3 *wt* and PNT3^3A^. We spectrophotometrically measured the red shift (from 497 nm to 515 nm) in the absorbance maximum of the CR spectrum of each sample following a four- or seven-days incubation at 37 °C. Results shown in **Figure 5** indicate that all the variants promote a shift in the CR spectrum whose amplitude increases with incubation time, suggesting that all the variants are able to progressively form amyloid-like fibrils or at least structures able to bind CR. The previous findings that pointed out that the HeV PNT3^3A^ variant has a reduced ability to bind CR compared to HeV PNT3 *wt* [25] were confirmed here (**Fig. 5**). Two of the three single alanine variants (PNT3^A1^, PNT3^A3^) show an intermediate behavior between HeV PNT3 *wt* and HeV PNT3^3A^, but without significant differences with either the *wt* or the PNT3^3A^ variant, while HeV PNT3^A2^ displays an ability to bind CR significantly higher than that of HeV PNT3^3A^ and similar to that of HeV PNT3 *wt* (**Fig 5**). These results therefore suggest that the central tyrosine in the motif could be less relevant to the fibrillation process. Notably, the NiV PNT3 variant shows a significantly decreased ability to bind CR compared to PNT3 *wt*, similar to that of the triple alanine variant. Thus, as already observed for the aggregation propensity, the reduced CR binding ability of NiV PNT3 likely results from its amino acid context rather than from the fact that it lacks a tyrosine in the motif.

**Figure 5.**
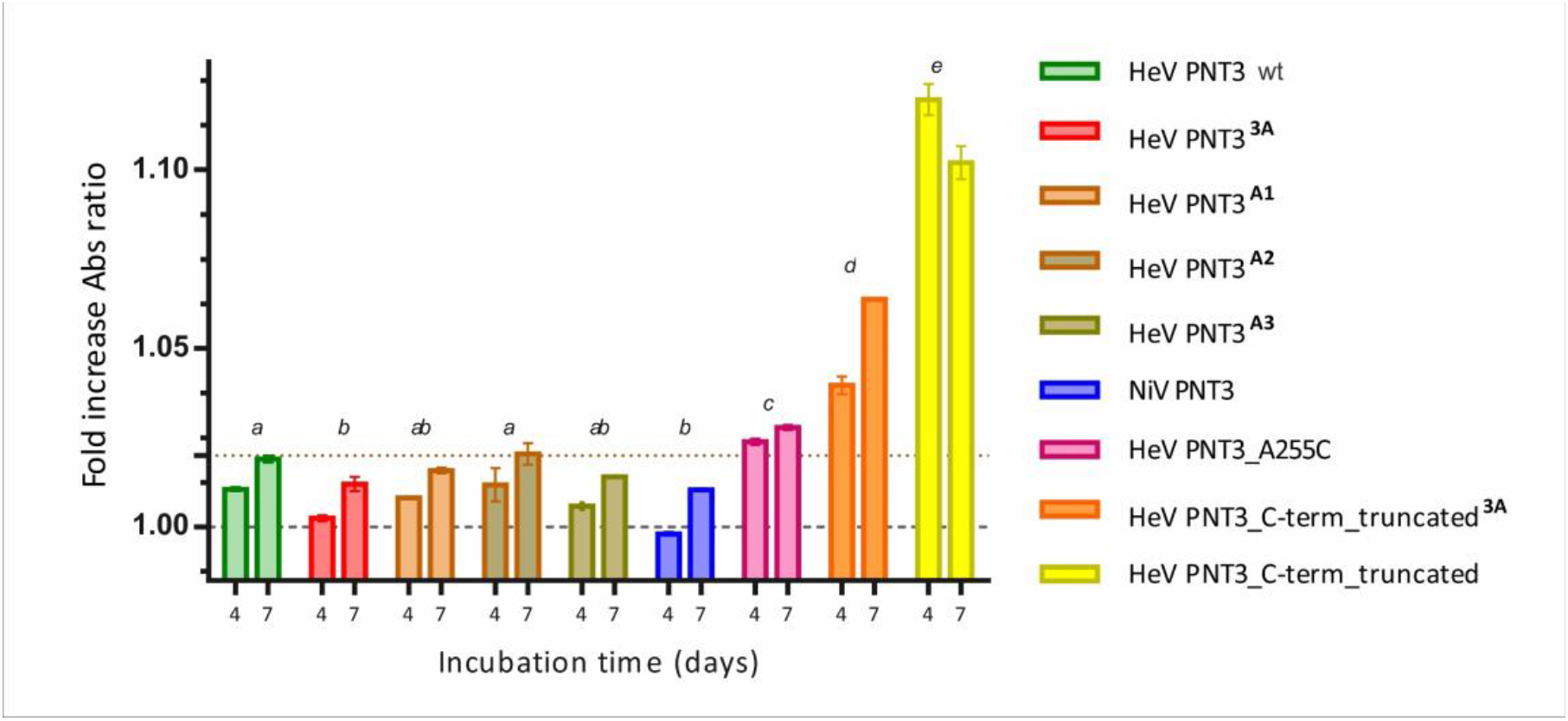
Congo Red binding assay of the set of PNT3 variants. The ability to bind CR is represented by the fold increase in the ratio between the absorbance at 515 and at 497 nm, with respect to a sample containing CR alone, of PNT3 samples at 20 μM after 4 and 7 days of incubation at 37 °C. The error bar corresponds to the standard deviation, with n = 3. Different letters indicate statistically significant differences (p < 0.05) (Two-way ANOVA test; *ab* means lack of statistically significant differences with respect to *a* or *b*).

#### 2.3.3. Propensity and time-dependance of fibrillation of the PNT3 EYYY motif variants using negative-staining transmission electron microscopy (ns-TEM)

To directly document fibril formation by the PNT3 EYYY variants as a function of time and to obtain orthogonal experimental evidence corroborating the CR binding assay results, we next carried out ns-TEM studies. These analyses were performed for each of the variants at 0, 24 and 96 h of incubation at 37 °C (**Fig. 6**). As shown the **Figure 6**, although HeV PNT3 *wt* in the selected conditions does not form fibrils at time 0, after 24 h of incubation it forms short fibrils, evolving to long fibrils after 96 h. In line with the CR binding results, ns-TEM studies confirmed that all PNT3 EYYY variants, including NiV PNT3, are able to form amyloid-like fibrils but with a significantly decreased ability compared to the *wt*. Specifically, after 24 h short fibrils can be observed in most variants, except for HeV PNT3^3A^. In an attempt at identifying possible significant differences among the EYYY variants in spite of their overall similar behavior, we performed a comprehensive analysis of the number and length of fibrils detected for each of them (**Fig. 7**). This analysis revealed no significant differences in the length of the fibrils obtained after 96 hours of incubation among the 3 variants with a single alanine (PNT3^A1^, PNT3^A2^ and PNT3^A3^). However, a slight, though significant, increase is observed in the PNT3^A3^ variant in the number of fibrils found *per* picture compared to the other two single alanine variants (**Fig. 7**). This slightly higher number of short fibrils observed for the PNT3^A3^ variant may indicate that the last tyrosine in the motif contributes less to the nucleation process compared to the two other tyrosines. Remarkably, the same analysis showed that the NiV PNT3 variant forms significantly longer fibrils compared to the PNT3^A1^, PNT3^A2^ and PNT3^A3^ variants, therefore confirming that other sequence attributes, beyond the amyloidogenic motif, contribute to the fibrillation process. In other words, fibril formation would rely not only on stabilizing contacts mediated by the E(H/Y)YY motif but also on additional stabilizing interactions established by other protein regions.

**Figure 6.**
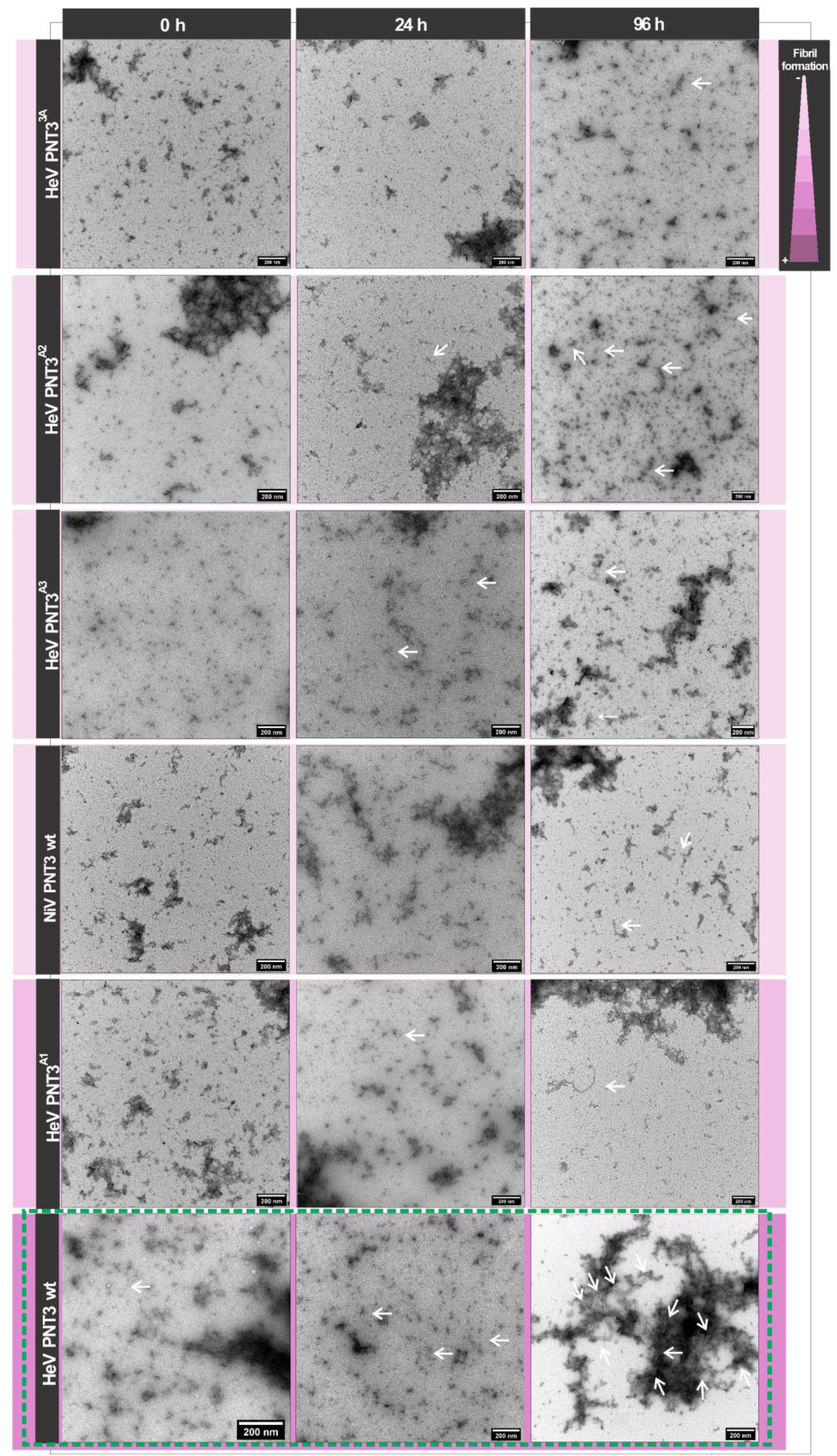
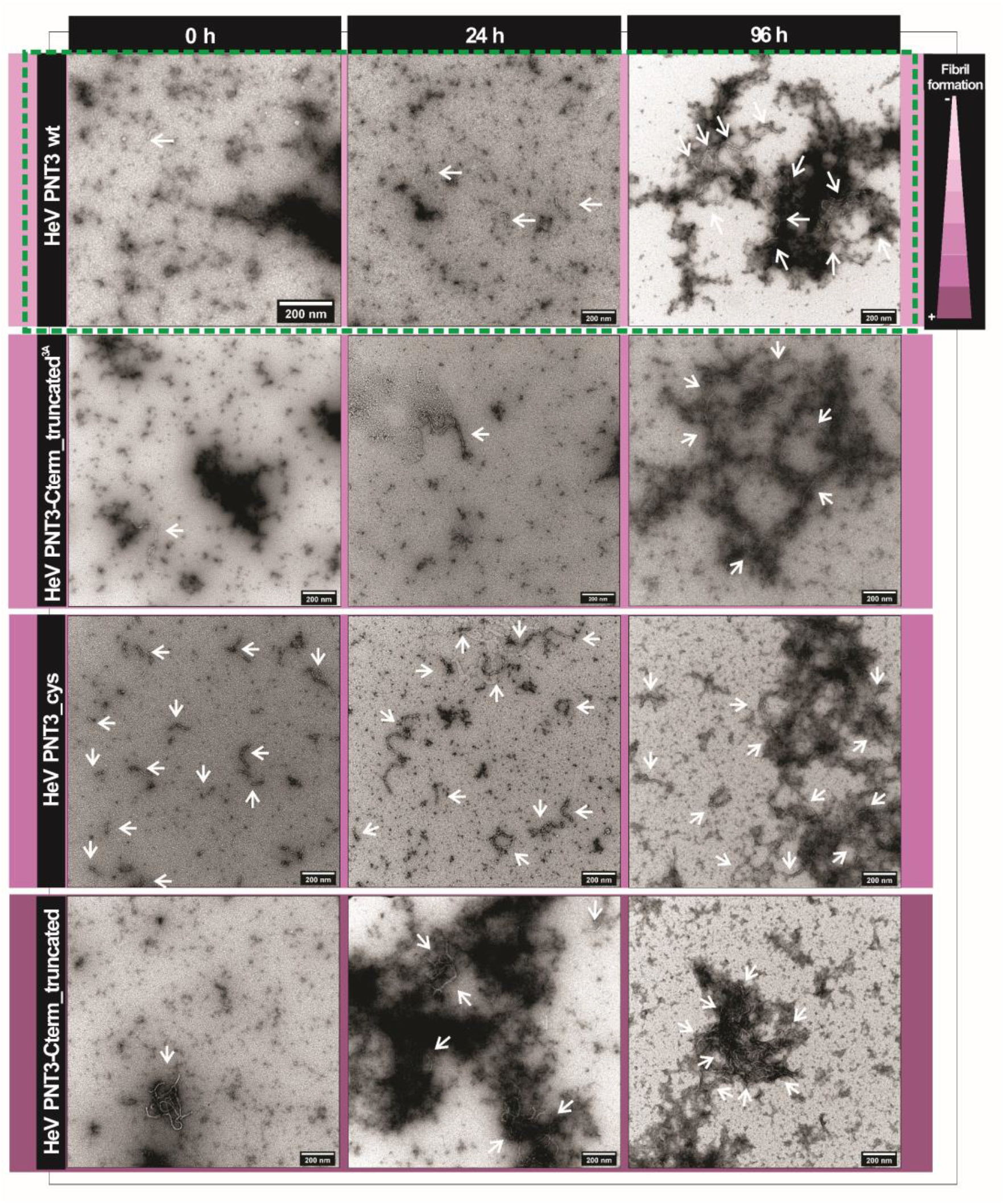
Fibril formation as a function of time by ns-TEM. Ns-TEM analysis of PNT3 variants (200 µM) at time zero and after 24 h or 96 h of incubation at 37 °C. Note that in all cases, samples were diluted to 40 µM prior to being deposited on the grid. White arrows indicate fibrils. The purple gradient represents the propensity to form fibrils.

**Figure 7.**
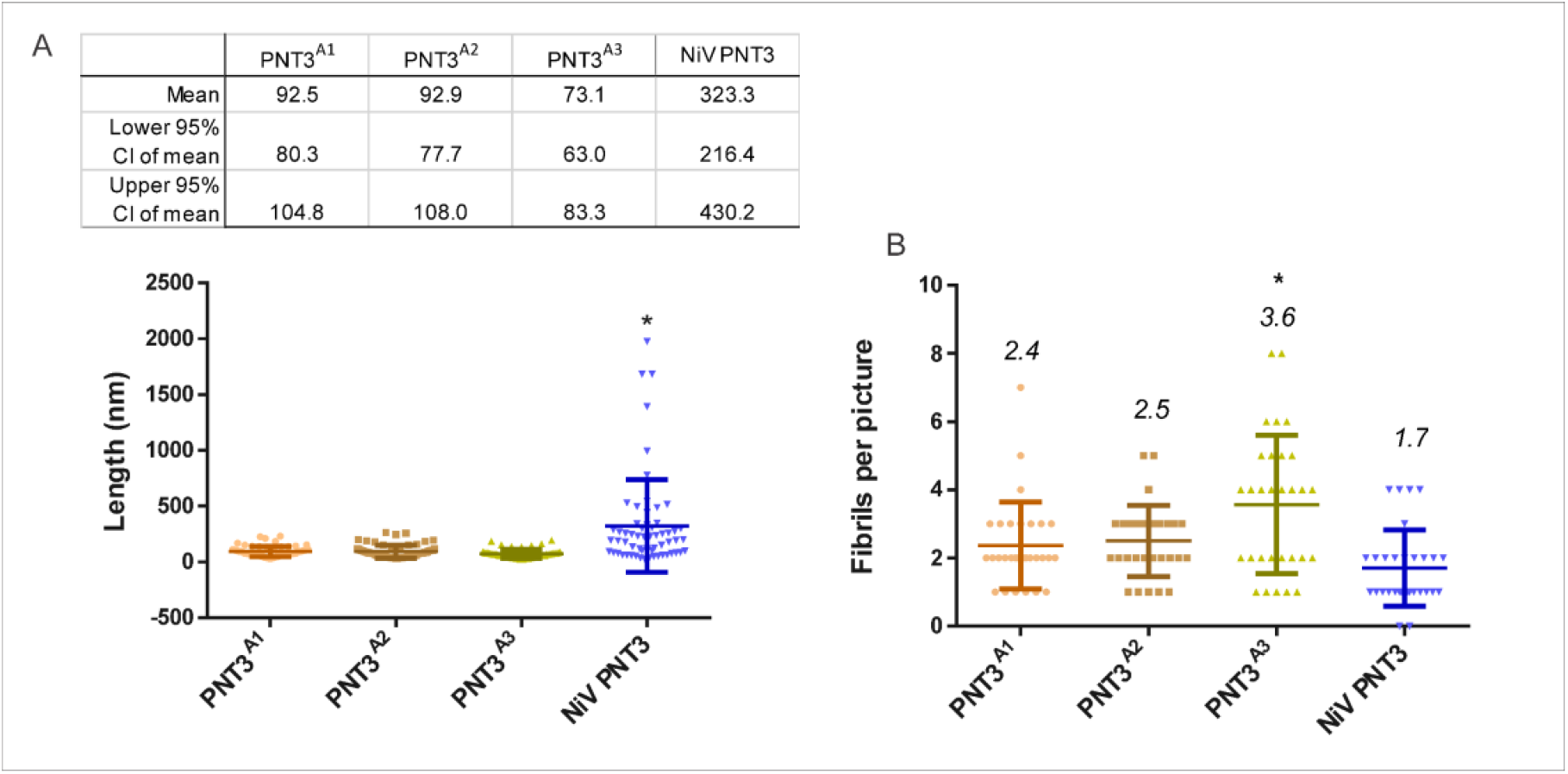
Fibril length of EYYY variants. **A**. For each variant, statistics on fibril length was obtained from analysis of the contour length of 60 fibrils after 96 h of incubation at 37 °C. The upper table shows the main statistic results. **B**. Number of fibrils detected *per* picture for each variant. Upper number indicates the mean value of each variant. The analysis was done using the ImageJ software. (*) indicates statistically significant differences (One-way Anova test, p-value < .05)

Altogether, CR binding assays and ns-TEM enabled documenting slight differences among the PNT3^A1^, PNT3^A2^ and PNT3^A3^ variants. Although other regions beyond the amyloidogenic motif were found to contribute to the fibrillation process, results strongly suggest that the absence of a single tyrosine in the EYYY motif leads to a significant decrease in the ability to form fibrils irrespective of the position. In light of the finding that the PNT3^A1^, PNT3^A2^ and PNT3^A3^ variants are able to form short fibrils, whose length does not increase under the experimental conditions herein used, we can assume that the three tyrosines mainly play a role in the elongation phase. The present results support the involvement of tyrosines in the formation of amyloid-like fibrils, with π-π stacking and H-bonding interactions between tyrosines likely allowing to form cross-β like architectures, as previously suggested [35].

### 2.4. Impact of the HeV PNT3 C-terminal region in fibrillation abilities

#### 2.4.1. Aggregation propensity of C-terminally truncated PNT3 variants

Taking into account the results presented above that lend support to a scenario where other motifs and/or sequence attributes beyond the amyloidogenic motif could be involved in the fibrillation process, we decided to investigate the impact of the PNT3 C-terminal region. We first studied the aggregation propensity of both PNT3 C-terminally truncated variants. **Figure 4A** shows that the PNT3 C-term truncated variant has the lowest PEG_1/2_ value, indicating a strikingly decreased relative solubility compared to all the full-length PNT3 variants. Notably, there are no significant differences in the PEG_1/2_ values between the PNT3 C-term truncated and its triple alanine mutant (PNT3^3A^ C-term truncated) (**Fig. 4C**). These results indicate that removal of the C-terminal region has a strong impact on the aggregation propensity, with this effect being insensitive to the sequence context of the EYYY motif.

#### 2.4.2. CR binding ability of C-terminally truncated PNT3 variants

Motivated by the results obtained by the PEG solubility assays pointing to a much higher aggregation propensity of both truncated variants, we next carried out CR binding assays. As shown in **Figure 5**, the HeV PNT3 C-terminal truncated variant displays a significantly increased ability to bind CR compared to full-length HeV PNT3 *wt*. In striking contrast with PEG solubility assays that detected no differences in terms of aggregation propensities between the two truncated variants, CR binding assays revealed significant differences between the two variants. In particular, the PNT3^3A^ truncated variant has a much-decreased ability to bind CR with respect to the truncated variant bearing a native EYYY motif, hence displaying an intermediate behavior between the full-length and truncated HeV PNT3 *wt* (**Fig. 5**). Therefore, it can be concluded that the C-terminal region, far from being inert, negatively affects the ability of the protein to form CR-binding structures. Albeit the PNT3^3A^ truncated variant binds more CR than the *wt* variant, the contribution of the EYYY motif is evident when the two truncated variants are compared. These results, however suggest that the EYYY motif would have only a marginal role in driving the formation of CR-binding structures in the context of C-terminally truncated form.

#### 2.4.3. Propensity and kinetics of fibrillation of C-terminally truncated PNT3 variants using ns-TEM studies

In order to ascertain whether the increased binding to CR and decreased solubility of the truncated variants is actually reflected in a higher fibrillation ability, we analyzed them by NS-TEM studies. In line with expectations, **Figure 6** clearly shows a significantly higher abundance of fibrils, as well as an increased fibril length, in the PNT3 C-terminal truncated variant at short incubation times, thus confirming its increased fibrillation potential. Notably, the PNT3^3A^ truncated variant displays a decreased fibrillation ability compared to its *wt* counterpart, similar to the full-length HeV PNT3 *wt* (**Fig. 6**). These findings suggest that the removal of the C-terminal region results in an acceleration of the fibrillation rate, reflecting an interaction between this region and the rest of the sequence that negatively affects the kinetics of fibril formation. Remarkably, similar results were previously documented in the case of the aggregation of α-synuclein (α-syn), an extensively characterized protein associated with neurodegeneration and whose transition from a soluble to a fibrillar form is thought to contribute to pathogenesis [36]. Compelling experimental indicates that C-terminal truncation of α-syn promotes *in vitro* oligomer and fibril formation (see [37] and references therein cited). The middle region of α-syn, referred to as “non-amyloid component” (NAC) domain, forms the core of α-syn filaments. The C-terminal region of α-syn can adopt conformations in which the C-terminus contacts the hydrophobic NAC domain thus shielding it from pathological templating interactions [37]. The negative charge of the C-terminal region has been proposed to contribute to this self-chaperoning activity *via* the establishment of electrostatic interactions [37].

In an attempt at rationalizing the observed self-inhibitory effect of the C-terminal region of PNT3 on fibril formation, we analyzed the charge distribution within the PNT3 sequence (**Supplementary Fig. S4** and **Table S2**). Although full-length PNT3, and its constituent N-terminal and C-terminal regions fall in very close positions in the phase diagram plot, the C-terminal region has a higher fraction of negatively charged residues compared to the N-terminal region (**Supplementary Fig. S4** and **Table S2**). In addition, the full-length form of PNT3 and the truncated variant strongly differ in their net charge at pH 7.0. Taking into account the strong impact of pH on PNT3 fibrillation, where a decrease to pH 6.5 strongly promotes fibrillation and leads to a behavior similar to that of the truncated PNT3 variant at pH 7.2, it is conceivable that the results obtained with the truncated variant could be, at least partly, accounted for by electrostatics, as in the case of α-syn. A plausible alternative scenario for the self-inhibitory effect of the C-terminal region of PNT3 on fibril formation could be the following: the disordered region downstream the fibril core may hamper fibril formation by slowing the disorder-to-order transition expected to take place in the core of the fibril, through either a purely entropic effect or through a combination of enthalpy and entropy as already documented in the case of fuzzy appendages adjacent to molecular recognition elements (for examples see [38,39]).

Altogether, these findings advocate for a key role of the C-terminal region in regulating the fibrillation properties of PNT3. In particular, the C-terminal region may act either as a molecular shield, as in the case of α-syn [37], or by slowing down the rate of folding of the core of the fibrils, with this property, irrespective of the underlying mechanisms, being also possibly relevant to biological function in *vivo*. Definite answers on the precise molecular mechanisms and on the possible biological relevance await future studies.

### 2.5. Impact of a cysteine in the HeV PNT3 sequence on fibrillation abilities

With the aim of elucidating the possible impact of a disulfide bridge-mediated PNT3 dimerization on its fibrillation abilities, we first studied the ability of the HeV PNT3 variant bearing a cysteine residue (PNT3_Cys) to bind CR. As shown in **Figure 5** this variant shows a significant increased ability to bind CR compared to HeV PNT3 *wt*. Subsequently, the ability of this variant to form fibrils was assessed by ns-TEM in the same conditions used for PNT3 *wt*. **Figure 6** shows that the PNT3_Cys variant is able to form fibrils even at time 0, indicating a higher fibrillation propensity compared to HeV PNT3 *wt*. Notably, the PNT3_Cys was the unique variant displaying an enrichment in shortened fibrils (**Fig. 6**). These findings suggest that the presence of one cysteine in the sequence mainly impacts the nucleation phase. We reasoned that the peculiar fibrillation behavior of this variant could result from disulfide bridge-mediated protein dimerization. However, the addition of DTT was found to exert a negligible impact, as judged from the presence at time 0 of very short fibrils and of long fibrils at 96 hours (**Supplementary Fig. S5**), resulting in an intermediate behavior between PNT3 *wt* and PNT3_cys under non-reducing conditions. Thus, the peculiar behavior of this variant cannot be ascribed to protein dimerization and rather stems from other intrinsic properties that remain to be elucidated.

### 2.6. Characterization of the aggregation process by Taylor Dispersion Analysis

We next sought at shedding light onto the fibrillation kinetics using Taylor Dispersion Analysis (TDA) [40]. TDA is a new technique in the field of protein aggregation that has the notable advantages of being able to (i) capture intermediate species, (ii) quantify early- and late-stage aggregates, and (iii) provide both kinetic and equilibrium constants. Besides, TDA is not dominated by aggregates (as opposite to scattering techniques), and is thus ideally suited to study the molecular mechanisms of protein fibrillation. Recently, this technique successfully allowed obtaining a complete quantitative picture of the aggregation process of both Aβ(1-40) and Aβ(1-42), including the size of the oligomers and protofibrils, the kinetics of monomer consumption, and the quantification of different early- and late-formed aggregated species [40-42].

In light of the fibrillation properties of all the PNT3 variants herein investigated, as revealed by the ensemble of studies described above, we decided to focus on the two variants with the most extreme phenotype, namely the variant with the highest fibrillation propensity, *i.e*., HeV PNT3_C-term_truncated, and the least fibrillogenic variant, *i.e*., PNT3^3A^. **Figure 8A** and **C** show a three-dimensional overview of the obtained taylorgrams during the aggregation process of the two selected variants. In the case of the truncated variant, two major species were detected over time: the monomer (hydrodynamic radius, *R*_*h*_, of ∼3 nm) (**Fig. 8B**) and large aggregates with an *R*_*h*_ ≥ 400 nm (see spikes at the beginning of the run in **Fig. 8A**). **Figure 8B** shows that the *R*_*h*_ value of the soluble fraction increases with time, reaching about the double of its initial value, when this species is present in low proportions. **Figure 8B** also shows that the monomeric population slowly decreases with time. The decrease of the peak area (*Y*) of the monomeric population could be fitted using a 1^st^ order exponential decay:

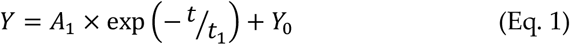

where *Y*_*0*_= 6.13 ± 0.20; *A*_*1*_= 7.63 ± 0.34 and *t*_*1*_= 12.06 ± 1.32 h. This resulted in a good quality fit as judged from the R^2^ value of 0.9394. From the fit, the kinetics of aggregation could be deduced, with a characteristic aggregation time of about *t*_*1*_= 12 h. From **Figure 8B**, the evolution in the spikes area gives an estimation of the quantity of large aggregated species entering the capillary. The proportion of these aggregated species increases with time until about 48 h, before decreasing because their size becomes too large to enter the capillary or because of precipitation in the sample vial. The presence of spikes at very short times of incubation suggests a fast aggregation process of the truncated form, while the absence of significant intermediate species suggests that the monomers add to the already present aggregates and elongate them.

**Figure 8.**
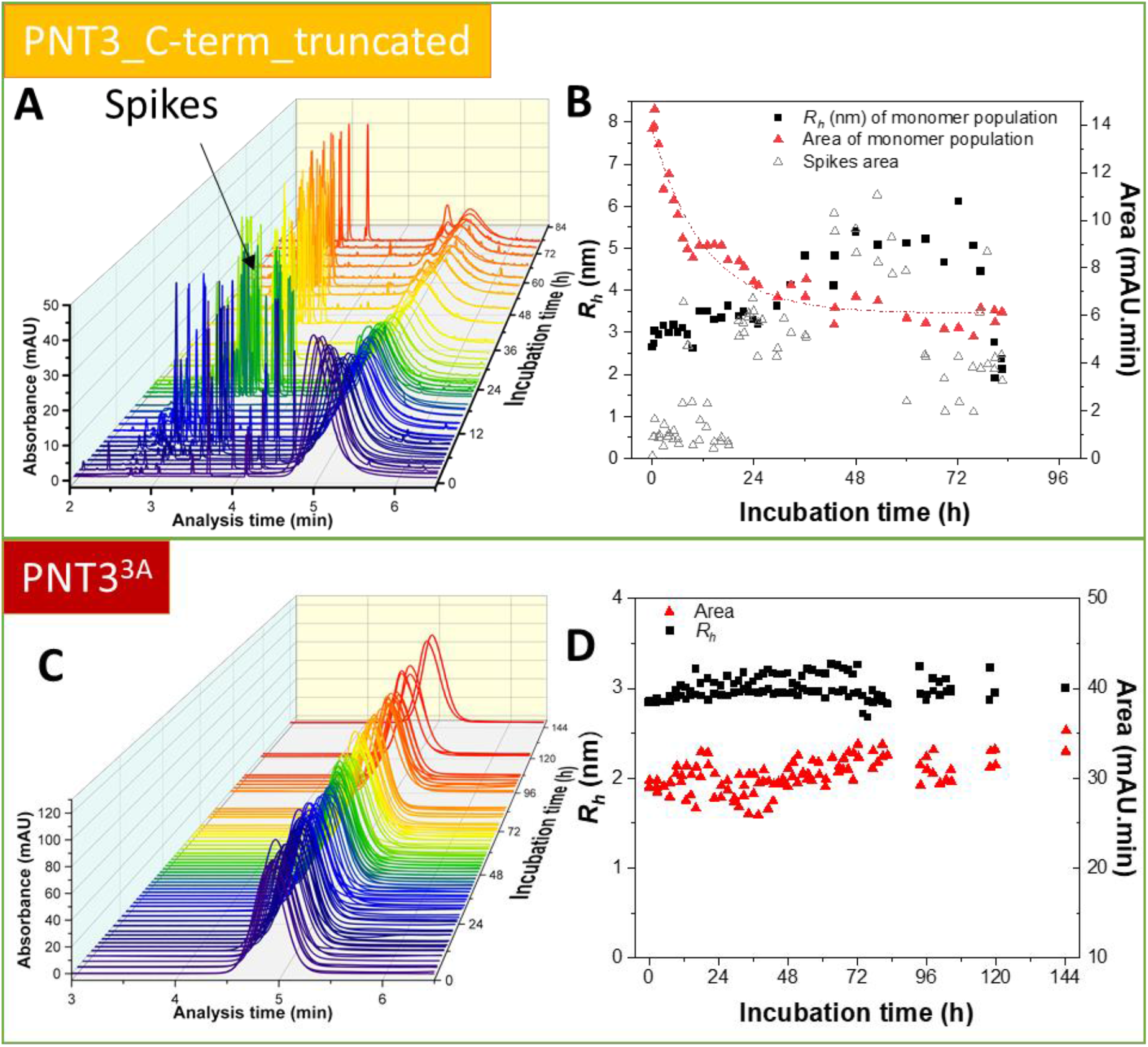
Kinetics of fibril formation by Taylor Dispersion Analysis (TDA). Three-dimensional overview of the obtained taylorgrams during the aggregation process of HeV PNT3-Cterm_truncated (**A**) and HeV PNT3^3A^ (**C**) at different incubation times. Analyses were performed using a protein concentration of 200 μM in 50 mM phosphate buffer, pH 7.2 at 37 °C. Peak area and hydrodynamic radius (*R*_*h*_) evolution of the monomeric species and of the spikes area during the aggregation process of HeV PNT3-Cterm_truncated (**B**) and HeV PNT3^3A^ (**D**).

By contrast, and in agreement with its dramatically reduced fibrillogenic abilities as unveiled by the other approaches herein used, the PNT3^3A^ variant does not show any changes during the incubation time, with both *R*_*h*_ and peak area remaining constant in this period (**Fig. 8C,D**).

In conclusion, the data support a much higher fibrillation ability for the truncated variant as corroborated by the presence of spikes in the elution profiles and by the fast consumption of the monomeric species as compared to PNT3^3A^ where no evolution in size nor in area was observed. The aggregation process of the truncated variant follows a first order kinetics, without significant formation of intermediate species between the monomeric and the fibrillar species. In PNT3^3A^ the fibrillation process could not be detected, indicating that the fibrils observed by ns-TEM represent a very poorly populated species within the system.

## 3. Conclusions

This study constitutes a comprehensively analysis of the molecular basis of the fibrillation process of a small region (PNT3) within the N-terminal intrinsically disordered domain shared by the HeV P/V/W proteins. Biochemical and biophysical characterization of a set of HeV PNT3 variants bearing alanine substitutions in the amyloidogenic EYYY motif, along with the characterization of the corresponding PNT3 region from the cognate NiV, revealed that each of the three tyrosines in the motif are required for the elongation step of the fibrillation process. Remarkably, the present study also unveiled a role for the C-terminal domain of PNT3 in self-inhibition of fibrillation, possibly reminiscent of the α-synuclein fibrillation model, and of potential biological significance. Noteworthy, in light of the observation that amyloid-like fibrils form not only *in vitro* but also the cellular context, it is tempting to hypothesize that the amyloidogenicity of V/W proteins, which both encompass the PNT3 region, could be correlated with the pathogenic (and even encephalitogenic) properties of Henipaviruses. Therefore, the PNT3 variants that we have herein generated constitute valuable tools to further explore the functional impact of V/W fibrillation in transfected and infected cells. The present results therefore set the stage for further investigations aimed at illuminating the mechanisms underlying the disease as a preliminary step towards the rational design of antivirals.

## 4. Materials and Methods

### 4.1. Generation of the constructs

The pDEST17OI/PNT3 and pDEST17OI/PNT3^3A^ expression plasmids, driving the expression of a hexahistidine tagged form of the protein of interest, have already been described [25]. For the construction of expression plasmids encoding the PNT3 variants bearing single alanine substitutions (PNT3^A1^, PNT3^A2^, PNT3^A3^), the pDEST17OI/PNT3 construct was used as template in two separate PCR amplifications using either primers attB1 and specific R_ala*N*-PNT3 (PCR1), or primers F_ala*N*-PNT3 and attB2 (PCR2), where *N* varies from 1 to 3 (see **Supplementary Table 1**). After DpnI treatment, 1 µl of PCR1 and 1 µl of PCR2 were used as overlapping megaprimers along with primers attB1 and attB2 in a third PCR. After purification, the third PCR product was inserted into the pDEST17OI bacterial expression vector using the Gateway® technology (Invitrogen). This vector allows expression of the recombinant protein under the control of the T7 promoter. The resulting protein is preceded by a stretch of 22 vector-encoded residues (MSYYHHHHHHLESTSLYKKAGF) encompassing a hexahistidine tag. The DNA fragment encoding NiV PNT3 (*i.e*., residues 200-314 of the NiV P/V/W protein) was PCR-amplified using the pDEST17OI/NiV W construct as template [21] and primers NiV PNT3-AttB1 and NiV PNT3-AttB2. After DpnI treatment, the resulting amplicon was cloned into pDEST17OI as described above.

The expression construct encoding the truncated HeV PNT3 variant (PNT3_C-term_truncated) was generated by PCR using pDEST17OI/PNT3 [25] as template and attB1 and Trunc_PNT3_B2 as primers. After DpnI treatment, the resulting amplicon was cloned in pDEST17OI. The same procedure was used to obtain the HeV PNT3 truncated variant bearing the triple alanine substitution (PNT3^3A^_C-term_truncated) except that the pDEST17OI/PNT3^3A^ construct [25] was used as template.

The construct encoding the HeV PNT3 variant bearing a cysteine (PNT3_Cys) was obtained using the pDEST17OI/PNT3 construct [25] as template in two separate PCR amplifications using either primers attB1 and specific PNT3_C255_R (PCR1), or primers PNT3_C255_F and attB2 (PCR2). After DpnI treatment, 1 µl of PCR1 and 1 µl of PCR2 were used as overlapping megaprimers along with primers attB1 and attB2 in a third PCR. After purification, the third PCR product was inserted into pDEST17. The list and sequence of primers used to generate the above-described constructs is provided in **Supplementary Table S1**. Primers were purchased from Eurofins Genomics. All the constructs were verified by DNA sequencing (Eurofins Genomics) and found to conform to expectations.

### 4.2. Proteins expression and purification

The *E. coli* strain T7pRos was used for the expression of all the recombinant proteins upon transformation of bacterial cells with each of the bacterial expression plasmids described above. Cultures were grown over-night to saturation in LB medium containing 100 µg mL^-1^ ampicillin and 34 µg mL^-1^ chloramphenicol. An aliquot of the overnight culture was diluted 1/20 into 1 L of TB medium and grown at 37 °C with shaking at 200 rpm. When the optical density at 600 nm (OD_600_) reached 0.5-0.8, isopropyl β-D-thiogalactopyanoside (IPTG) was added to a final concentration of 0.5 mM, and the cells were grown at 35 °C overnight. The induced cells were harvested, washed and collected by centrifugation (5,000 g, 15 min). The resulting pellets were resuspended in buffer A (50 mM Tris/HCl pH 7.5, 1 M NaCl, 20 mM imidazole) containing 6 M guanidium hydrochloride (GDN). The suspension was sonicated to disrupt the cells (using a 750 W sonicator and 3 cycles of 30 s each at 45 % power output) and then centrifuged at 14,000 g for 30 min at 20 °C. The supernatant was first purified by Immobilized Metal Affinity Chromatography (IMAC) by loading the clarified lysate onto a 5 mL Nickel column (GE Healthcare) pre-equilibrated in buffer A. The affinity resin was washed with 20 column volumes (CV) of buffer A. Proteins were eluted with ∼3 CV of buffer A supplemented with 250 mM imidazole. The fractions eluted from the Nickel column were pooled and concentrated in the presence of 6 M GDN up to 1 mM using Centricon concentrators, and the proteins were then frozen at -20 °C. All PNT3 variants were subsequently subjected to SEC, where the SEC column was equilibrated with buffer B (Sodium Phosphate 50 mM pH 7.2 100 mM NaCl, 5 mM EDTA). The fractions from SEC, were pooled, supplemented with 6M GDN and concentrated (up to ∼750 µM) and stored at -20 °C. In the case of the PNT3 cysteine variant, the GDN-containing sample was also supplemented with 10 mM DTT. Prior to each subsequent analysis, the samples were loaded onto a Sephadex G-25 medium column (GE Healthcare) to exchange the buffer. The proteins were eluted from G-25 columns using Sodium Phosphate 50 mM buffer at a pH of 7.2 unless differently specified. IMAC and SEC were performed at room temperature (RT).

Protein concentrations were estimated using the theoretical absorption coefficients at 280 nm as obtained using the program ProtParam from the EXPASY server.

The purity of the final purified products was assessed by SDS-PAGE (**Fig. 3B**). The identity of all the purified PNT3 variants generated in this work was confirmed by mass spectrometry analysis of tryptic fragments obtained after digestion of the purified protein bands excised from SDS-polyacrylamide gels (data not shown). The excised bands were analyzed by the mass spectrometry facility of Marseille Proteomics in the same way as previously done for the PNT3 wt variant [25]. Briefly, proteins were reduced by incubation with 100 mM dithiothreitol (DTT) for 45 minutes at 56 °C and free cysteine residues were capped by incubation with 100 mM iodoacetamide for 30 h at 25 °C in the dark. Samples were digested with porcine trypsin (V5111, Promega) at 12,5 ng/µl in 25 mM NH_4_HCO_3_, for 18 h at 37 °C. Peptides were extracted from the gel with 60% vol/vol acetonitrile in 5% formic acid, dried under vacuum, and reconstituted in 5 µl of 50% vol/vol acetonitrile in 0.3% trifluoroacetic acid. Mass analyses of the tryptic fragments were performed on a MALDI-TOF-TOF Bruker Ultraflex III spectrometer (Bruker Daltonics, Wissembourg, France) controlled by the Flexcontrol 3.0 package (Build 51). This instrument was used at a maximum accelerating potential of 25 kV and was operated in reflectron mode and an m/z range from 600 to 4500 (RP_Wide range_Method) or 600 to 3400 (RP_Proteomics_2015_Method). The laser frequency was fixed to 200 Hz and approximately 1000-1500 shots by sample were cumulated. Five external standards (Peptide Calibration Standard, Bruker Daltonics) were used to calibrate each spectrum to a mass accuracy within 50 ppm. Peak picking was performed with Flexanalysis 3.0 software (Bruker) with an adapted analysis method. Parameters used were as follows: SNAP peak detection algorithm, S/N threshold fixed to 6 and a quality factor threshold of 30. One µl of sample was mixed with 1 µl of a saturated HCCA (α-cyano-4-hydroxycinnamic acid) solution in acetonitrile/0.3%TFA (1:1) then 1 µl was spotted on the target, dried and analyzed with the previously described method. Because of the high sequence similarity between the PNT3^A1,2,3^ variants, a MSMS analysis was necessary to unambiguously identify each of them.

### 4.3. PEG precipitation assay (relative solubility)

The relative solubility of each variant was evaluated at different PEG concentrations using an adaptation of the protocol recently described by Oeller et al. [32]. Briefly, PEG solutions from 0 to 30 % were prepared from 50 % PEG_6000_ stock solution. Then, an aliquot of corresponding protein from a stock (at 260 µM) was mixed with each PEG solutions to obtain a final concentration of 66 µM protein in a 100 µL final reaction volume. The assay was performed in 96-well plates sealed with aluminum plate sealers to prevent possible evaporation (Corning). Plates were incubated at 4 °C for 24 hours and centrifugated at maximum velocity (4600 x g) for 2 hours. Immediately after, 2 µL of the supernatant were pipetted to quantify the soluble protein concentration using a ND-1000 Nanodrop Spectrophotometer and theoretical absorption coefficients at 280 nm as obtained using the program ProtParam from the EXPASY server. Each condition was made in triplicate. The soluble fractions obtained were normalized and fitted to a sigmoid function to obtain the PEG_1/2_ value that reports on relative solubility (PRISM software). The error on the PEG_½_ and the quality of the fit were estimated by a 95 % confidence interval analysis (PRISM software).

### 4.4. Far-UV circular dichroism

CD spectra were measured using a Jasco 810 dichrograph, flushed with N_2_ and equipped with a Peltier thermoregulation system. Proteins were loaded into a 1-mm quartz cuvette at 0.06 mg/mL (in 10 mM phosphate buffer at pH 7.2) and spectra were recorded at 37 °C. The scanning speed was 20 nm min^-1^, with data pitch of 0.2 nm. Each spectrum is the average of ten acquisitions. The spectrum of buffer was subtracted from the protein spectrum. Spectra were smoothed using the ‘‘means-movement’’ smoothing procedure implemented in the Spectra Manager package. As already previously documented, a decrease in the signal spectrum was observed with increasing incubation time and ascribed to fibril formation [25]. Because the different variants have different fibrillation propensities, differences in spectra intensity might reflect differences in the fraction of fibrillar species (which are not detected by CD) rather than *bona fide* spectral differences. We therefore normalized spectra using the maximum negative value of intensity as a normalization factor.

Mean molar ellipticity values per residue (MRE) were calculated as

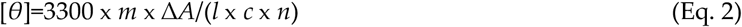

where *l* is the path length in cm, *n* is the number of residues, *m* is the molecular mass in Daltons and *c* is the concentration of the protein in mg mL^-1^.

### 4.5. Estimation of the hydrodynamic radius by SEC

The hydrodynamic radii (Stokes radii, R_S_) of the proteins were estimated by analytical SEC using a HiLoad 16/600 Superdex 75 pg column (Cytiva). Buffer B was used as elution buffer. Typically, 250 μL of purified protein at 11 mg mL^-1^ were injected.

The Stokes radii of proteins eluted from the SEC column were deduced from a calibration curve obtained using globular proteins of known R_S_ (Conalbumin: 36.4 Å, Carbonic Anhydrase: 23 Å, RNAse A: 16.4 Å Aprotinin:13.5 Å)

The R_S_ (in Å) of a natively folded (*Rs*^*NF*^), fully unfolded state in urea (*R*_*S*_^*U*^) and natively unfolded premolten globule (PMG) (*R*_*S*_^*PMG*^) protein with a molecular mass (*MM*) (in Daltons) were calculated according to [43]:

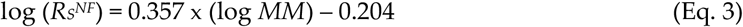

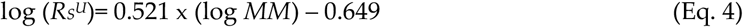

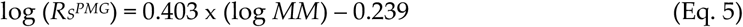

The *R*_*S*_ (in Å) of an IDP with *N* residues was also calculated according to [44] using the simple power-law model:

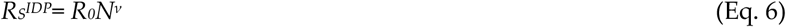

where R_0_ = 2.49 and ν = 0.509. The compaction index (*CI*) is expressed according to [45]:

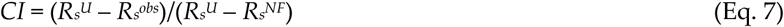

This parameter, which allows comparison between proteins of different lengths, in principle varies between 0 and 1, with 0 indicating minimal compaction and 1 maximal compaction.

### 4.6. Small-angle X-ray scattering (SAXS)

In order to ensure maximal monodispersity of the sample, SAXS studies were coupled to SEC. SEC-SAXS data were collected at SOLEIL (Gif-sur-Yvette, France) as described in **Table 3**. In both cases, the calibration was made with water. Sample from each PNT3 variant at 5 mg mL^-1^ in buffer B containing 6M GDN was injected onto an AdvanceBio SEC 2.7 µm (Agilent) SEC column. Elution was carried out in buffer C (50 mM sodium phosphate buffer at pH 7.2). Data reduction and frames subtraction were done with the beamline software FOXTROT. Gaussian decomposition was performed using the UltraScan solution modeler (US-SOMO) HPLC-SAXS module [46] or Chromixs (manual frames selection) [47] and the final deconvoluted scattering curves were submitted to the SHANUM program [48] to remove noisy, non-informative data at high angles.

**Table 3.** SEC-SAXS data acquisition parameters.

|  |  |
| --- | --- |
| Instrument | SOLEIL Synchrotron<br>(Gif-sur-Yvette, France)<br>Beamline Swing |
| X-rays wavelength (Å) | 1.033 |
| Energy (keV) | 12 |
| Detector type | Dectris EIGER 4M |
| Sample-to-detector distance (m) | 2.0 |
| q-range | 0.003 – 0.549 Å<sup>-1</sup> |
| Temperature (°C) | 20 |
| **Samples** |  |
| Concentration (mg mL<sup>-1</sup>) | 5 |
| Sample volume (μL) | 50 |
| Gel filtration column | AdvanceBio SEC 2.7 μm (Agilent) |
| Flow rate (mL min<sup>-1</sup>) | 0.3 |
| Buffer | 50 mM sodium phosphate pH 7.2 (buffer C) |

The data were analyzed using the ATSAS program package [48]. The radius of gyration (R_g_) and I(0) were estimated at low angles (*q.R*_*g*_ < 1.3) according to the Guinier approximation [49,50]:

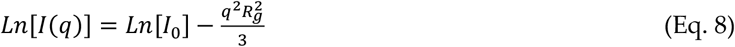

The pairwise distance distribution functions P(r), from which the *D*_*max*_ and the *R*_*g*_ were estimated, were calculated with the program GNOM [51] and manually adjusted until a good CorMap p-value (α >0.01) was obtained [51].

The theoretical *R*_*g*_ value (in Å) expected for various conformational states was calculated using Flory’s equation

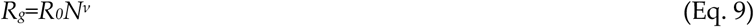

where *N* is the number of amino acid residues, *R*_*0*_ a constant and ν a scaling factor. For IDPs, *R*_*0*_ is 2.54 ± 0.01 and *ν* is 0.522 ± 0.01 [52], for chemically denatured (U) proteins *R*_*0*_ is 1.927 ± 0.27 and *ν* is 0.598 ± 0.028 [52], and for natively folded (NF) proteins *R*_*0*_= √(3/5) x 4.75 and *ν* = 0.29 [53].

As in the case of the *R*_*s*_, the *CI* allows comparing the degree of compaction of a given IDP, through comparison of the observed *R*_*g*_ to the reference values expected for a fully unfolded and a folded conformation of identical mass. The *CI* referred to the *R*_*g*_ can be calculated as follows [45]:

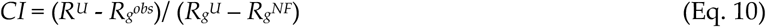

where *R*_*g*_^*obs*^ is the experimental value for a given protein, and *R*_*g*_^*U*^ and *R*_*g*_^*NF*^ are the reference values calculated for a fully unfolded (U) and natively folded (NF) form as described above. Akin the *R*_*S*_-based *CI*, this index increases with increasing compaction.

The overall conformation and the flexibility of the proteins was assessed with the dimensionless Kratky plot ((*q*R_g_)^2^ I(*<u>q</u>*)/I_0_ *vs q*R_g_) and the Krakty-Debye plot (*q*^2^I(*q*) *vs q*^2^).

SEC-SAXS data have been deposited in the Small Angle Scattering Biological Data Bank (SASBDB) [54] under codes SASDQB7, SASDQC7, SASDQD7, SASDQE7 and SASDQF7 for the set of data of PNT3 *wt*, PNT^3A^, NiV PNT3, PNT3_C-terminal truncated and PNT3^3A^_C-terminal truncated, respectively.

### 4.7. Congo red binding assays

Quantitative measurement of Congo Red (CR, Sigma-Aldrich) binding (CR shift assay) was carried out by using protein samples containing each PNT3 variant at 20 μM (in Buffer C) and 5 μM of CR in a final volume of 100 μL. The samples were then incubated at 37 °C for 4 or 7 days. The adsorption spectrum of the CR-containing samples was recorded using a PHERAstar FSX Microplate Reader (BMG LABTECH) in the 350-600 nm wavelength range. A solution of 5 μM CR in Buffer C without the protein was used as a control to normalize the analysis. Experiments were carried out in triplicate. Statistical analysis was done using Two-way ANOVA test implemented in the PRISM software.

### 4.8. Negative-staining transmission electron microscopy (TEM)

All the variants, at a concentration of 200 µM, were prepared and analyzed at different times to monitor their evolution (0, 24, 96 h). Incubation was carried out at 37 °C in Buffer C. Prior to each measurement, the samples were diluted to reach a final concentration of 40 µM. EM grids (carbon coated copper grids, 300 mesh, Agar Scientific) were exposed to plasma glow discharge for 20 seconds using GloQube (Quorum) (Current 15mA) in order to increase protein adhesion. Drops of 3.5 μL of the diluted protein solutions were deposited onto glow-discharged grids. After 1min incubation with the sample, the grids were washed three times with 50 µL of buffer C, once in 35 µL 1 % (w/v) Uranyl acetate solution (LauryLab-France) and then stained for 1 min in the latter solution. Excess of uranyl was blotted and grids were left to dry for 1 h at RT. Images were collected on Tecnai 120 Spirit TEM microscope (Thermo Fisher Scientific) operated at 120 kV using a Veleta 2Kx2K CCD camera (Olympus).

### 4.9. Kinetic protein aggregation study by Taylor Dispersion Analysis (TDA)

TDA was performed as already described [40] using an Agilent 7100 (Waldbronn, Germany) capillary electrophoresis system with bare fused silica capillaries (Polymicro Technologies) having 60 cm × 50 μm i.d. dimensions and a detection window at 51.5 cm. New capillaries were conditioned with the following flushes: 1 M NaOH for 30 min and ultrapure water for 30 min. Between each analysis, the capillaries were rinsed with Buffer C (2 min). Samples were injected hydrodynamically on the inlet end of the capillary (30 mbar, 6 s, injected volume is about 6.1 nL corresponding to less than 1% of the capillary volume to the detection point). Experiments were performed using a mobilization pressure of 100 mbar. The temperature of the capillary cartridge was set at 37 °C. The vial carrousel was thermostated using an external circulating water bath from Instrumat (France). The solutes were monitored by UV absorbance at 198 nm. The mobile phase was Buffer C. (viscosity at 37 °C is 7.54 × 10^−4^ Pa s). Samples obtained after the desalting column were diluted to reach 700 μL at 200 μM solution, and were immediately transferred to a vial and incubated at 37 °C in the capillary electrophoresis instrument’s carrousel. The aggregation was conducted by injecting the sample every 1 h. The total number of TDA runs for each sample was about 120. The taylorgrams were recorded with Agilent Chemstation software and then exported to Microsoft Excel for subsequent data processing. Data were fitted to a first order exponential decay according to equation 1. This resulted in a good quality fit as judged from the Reduced Chi-Sqr: 0.43723; R-Square (COD)= 0.93943, and Adj. R-Square= 0.93576. When necessary, the elution peaks were fitted with the sum of *n* Gaussian functions (in this work: *n* ≤ 3) as already described [40,41].

## Acknowledgements

We thank Aurélien Thureau (SOLEIL) and Anton Popov (ESRF) for their help in recording SEC-SAXS data. We thank both the ESRF and the SOLEIL synchrotrons for beamtime allocation. We are also grateful to Gerlind Sulzenbacher (AFMB lab) for efficiently managing the AFMB BAG. We thank Patrick Fourquet for mass spectrometry analyses done using the mass spectrometry facility of Marseille Proteomics (marseille-proteomique.univ-amu.fr), supported by IBISA (Infrastructures Biologie Santé et Agronomie), Plateforme Technologique Aix-Marseille, the Cancéropôle PACA, Région Sud-Alpes-Côte d’Azur, the Institut Paoli-Calmettes, the Centre de Recherche en Cancérologie de Marseille (CRCM), Fonds Européen de Développement Régional and Plan Cancer.

