## Supplementary Information for "Molecular determinants of fibrillation in a viral amyloidogenic domain from combined biochemical and biophysical studies"

**Table S1.** Nucleotide sequence of the primers used to generate the various constructs.

| Primer Name | Sequence (5'-3') | Name of the construct | Reference |
|---|---|---|---|
| HeV PNT3-AttB1 | GGGGACAAGTTTGTACAAAAAAGCAGGCTTCACCCCGACCGAAGAACCGCCG | PNT3-pDEST17 | Salladini, et al., 2021 |
| HeV PNT3-AttB2 | GGGGACCACTTTGTACAAGAAAGCTGGGTCTTATTAGCTATCTTTACGACGCACATC | | |
| attB1 | ACAAGTTTGTACAAAAAAGCAGGCT | | |
| attB2 | ACCACTTTGTACAAGAAAGCTGGGT | | |
| R_3ala-PNT3 | TCGCCACGACGGCCGCTACCGGCTGCCGCTTCCGGAATAACCGGCGGTTCTTC | $PNT3^{3A}$-pDEST17 | Salladini, et al., 2022 |
| F_3ala-PNT3 | AACCGCCGGTTATTCCGGAAGCGGCAGCCGGTAGCGGCCGTCGTGGCGATCTG | | |
| F_ala1_PNT3 | CGGAAGCGTATTATGGTAGCGGCCGTCGTGGCGA | $PNT3^{A1}$-pDEST17 | This work |
| R_ala1_PNT3 | GCTACCATAATACGCTTCCGGAATAACCGGCGGTT | | |
| F_ala2_PNT3 | CGGAATATGCGTATGGTAGCGGCCGTCGTGGCGA | $PNT3^{A2}$-pDEST17 | This work |
| R_ala2_PNT3 | GCTACCATACGCATATTCCGGAATAACCGGCGGTT | | |
| F_ala3_PNT3 | CGGAATATTATGCGGGTAGCGGCCGTCGTGGCGA | $PNT3^{A3}$-pDEST17 | This work |
| R_ala3_PNT3 | GCTACCCGCATAATATTCCGGAATAACCGGCGGTT | | |
| NiV PNT3-AttB1 | ACAAGTTTGTACAAAAAAGCAGGCTCCGATCCTGCAAAAGACTCTCC | NiV_PNT3-pDEST17 | This work |
| NiV PNT3-AttB2 | ACCACTTTGTACAAGAAAGCTGGGTTTATTATGAGTCCTTTGGACCGGCAC | | |
| Trunc_PNT3_B2 | ACCACTTTGTACAAGAAAGCTGGGTCTTATTAAAATTCATCTTCATATTCCAG | PNT3_C-term_truncated - pDEST17 | This work |
| PNT3_C255_F | CTGGAATATGAAGATGAATTTTGCAAAAGCAGCAGCGAAGTGGTG | PNT3_Cys-pDEST17 | This work |
| PNT3_C255_R | AAATTCATCTTCATATTCCAG | | |

**Table S2**. Analysis of physicochemical properties of full-length PNT3 *wt*, PNT3 truncated and C-terminal region as provided by CIDER (http://pappulab.wustl.edu/CIDER/) [1].

| | PNT3 *wt* | PNT3_trunc | C-terminal |
|---|---|---|---|
| **N** | 133 | 77 | 56 |
| **f-** | 0.22556 | 0.19481 | 0.26786 |
| **f+** | 0.09774 | 0.09091 | 0.10714 |
| **FCR** | 0.32331 | 0.28571 | 0.37500 |
| **NCPR** | -0.12782 | -0.10390 | -0.16071 |
| **Kappa** | 0.25641 | 0.36530 | 0.17898 |
| **Hydropathy** | 3.27594 | 3.25584 | 3.30357 |
| **Phase Plot Region** | 2 | 2 | 3 |
| **Phase Plot Annotation** | Boundary Region | Boundary region | Coils,Hairpins and Chimeras |
| **Net charge at pH 7.0** | -16.34 | -7.44 | |

**N:** Number of residues; **f-:** Fraction of negatively charged residues; **f+:** Fraction of positively charged residues; **FCR:** Fraction of charged residues; **NCPR:** Net charge per residue; **Kappa:** κ (charge patterning parameter); **Hydropathy:** The 0-9 scaled Kyte-Doolittle hydropathy score for the sequence (9 most hydrophobic, 0 least hydrophobic); **Phase Plot Region:** Location on the Das-Pappu phase plot this sequence falls; **Phase Plot Annotation:** Annotation associated with a specific region of the Das-Pappu phase plot.

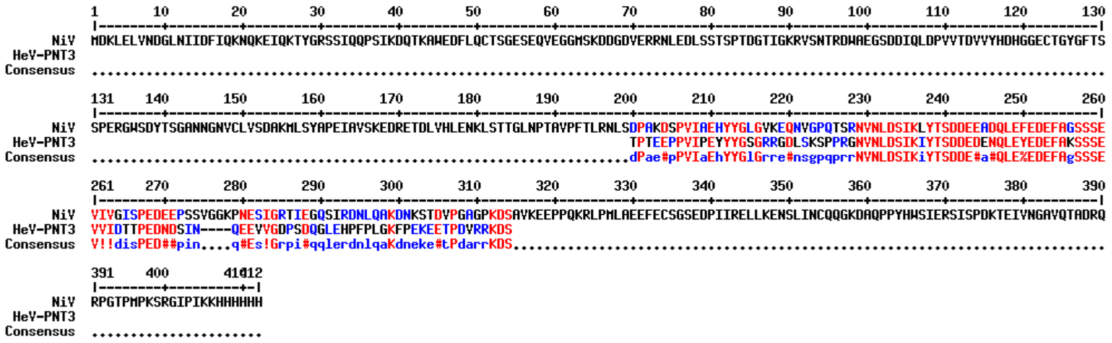


**Figure S1.** Alignment of NiV PNT3 and HeV PNT3 regions.

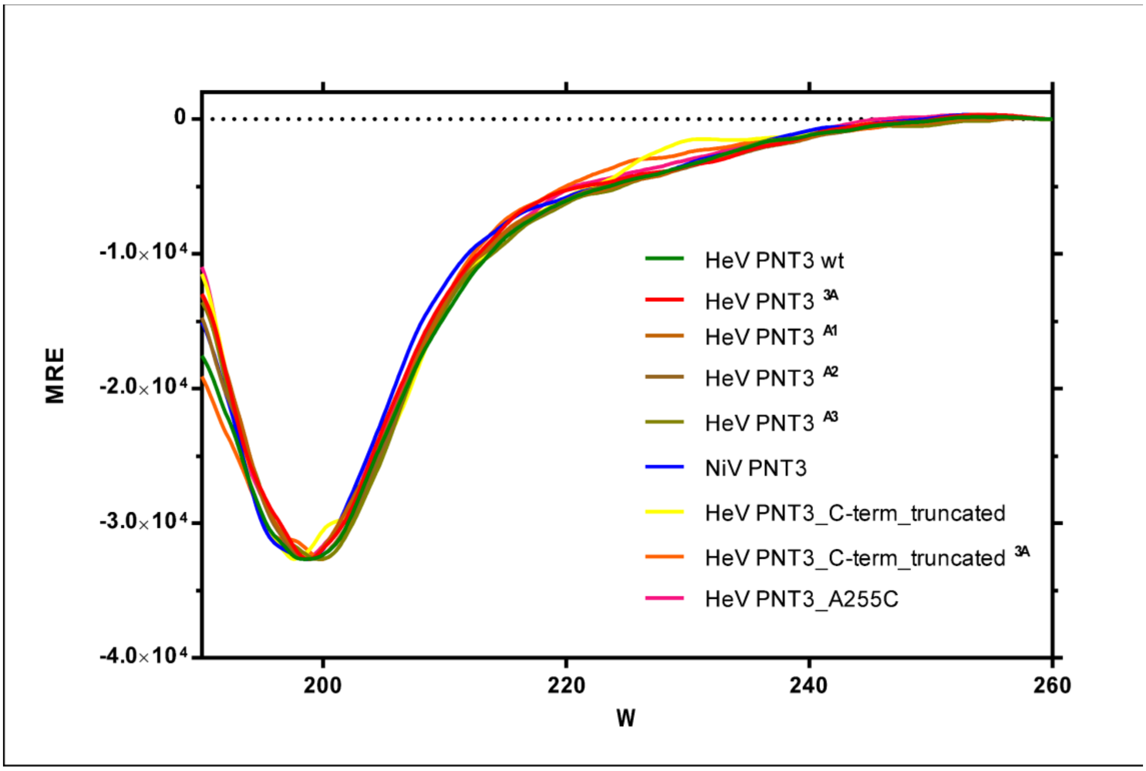


**Figure S2**. Far-UV circular dichroism (CD) studies of PNT3 variants. **A.** Spectra of PNT3 variants recorded in 10 mM sodium phosphate pH 7.2 at 37 °C. Protein concentration was 0.06 mg mL$^{-1}$ (4 μM). Spectra were recorded from a freshly purified PNT3 variant sample recorded immediately after elution from the SEC column. MRE (Θ) is expressed in deg cm$^2$ dmol$^{-1}$.

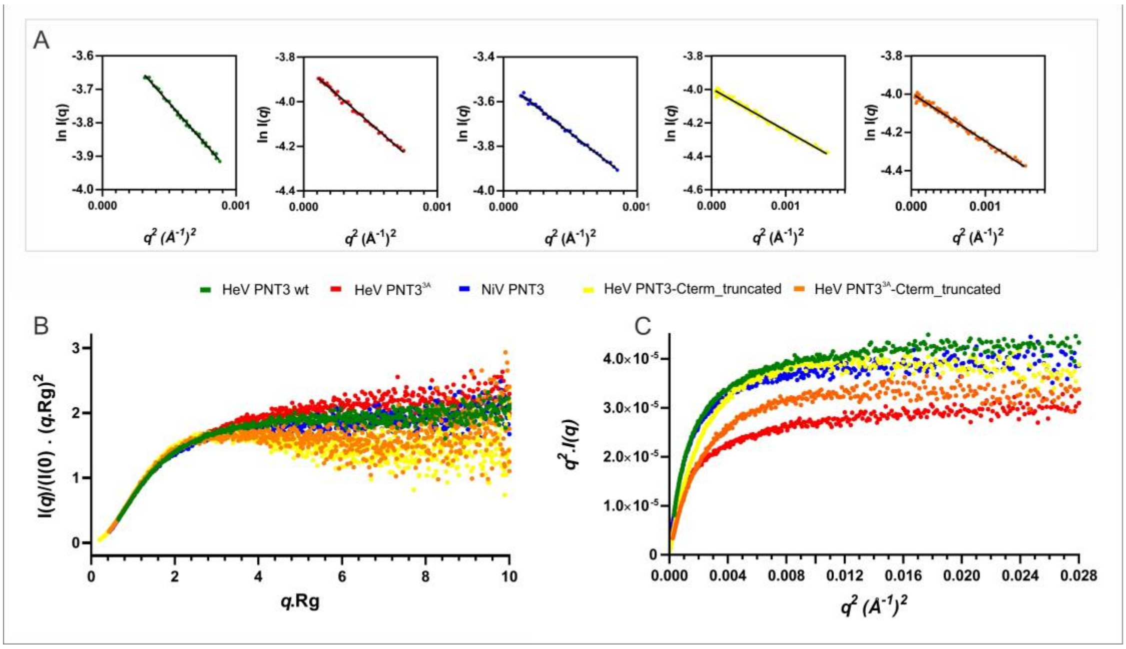


**Figure S3.** SEC-SAXS analysis of the monomeric form of PNT3 variants. **A.** Guinier plots of the experimental scattering curves used to determinate the radius of gyration ($R_g$) of th variants at low angles ($q_{max}$.Rg < 1.1)). **B**. Normalized Kratky plot. **C.** Kratky-Debye plot.

*MSYYHHHHHH LESTSLYKKA GF*TPTEEPPV IPEYYYGSGR RGDLSKSPPRGNVNLDSIKI YTSDDEDENQ
LEYEDEFAKS SSEVVIDTTP EDNDSINQEEVVGDPSDQGL EHPFPLGKFP EKEETPDVRR KDS

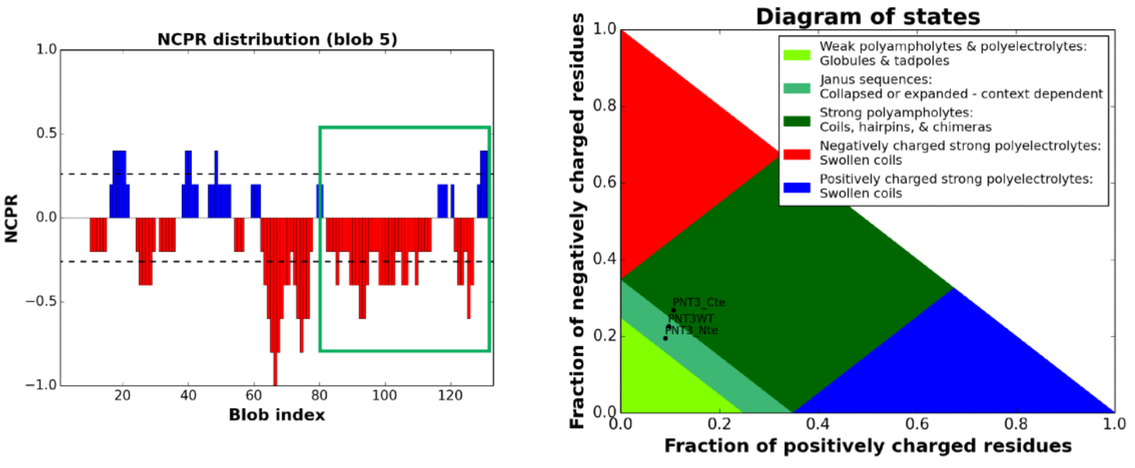


**Figure S4.** Analysis of charge distribution within the PNT3 sequence. The amino acid sequence of the recombinant protein is shown on the top, with vector-encoded residues shown in italic on a grey background, and residues removed in the PNT3 C-terminally truncated variant underlined. Left panel: net charge per residues (NCPR) as a function of residue number, with the C-terminal region framed. Right panel: phase diagram plot of full-length PNT3 *wt*, and of its N-terminal (Nte, aa 200-254) and C-terminal (Cte, aa 255-310) regions, as provided by CIDER (http://pappulab.wustl.edu/CIDER/) [1].

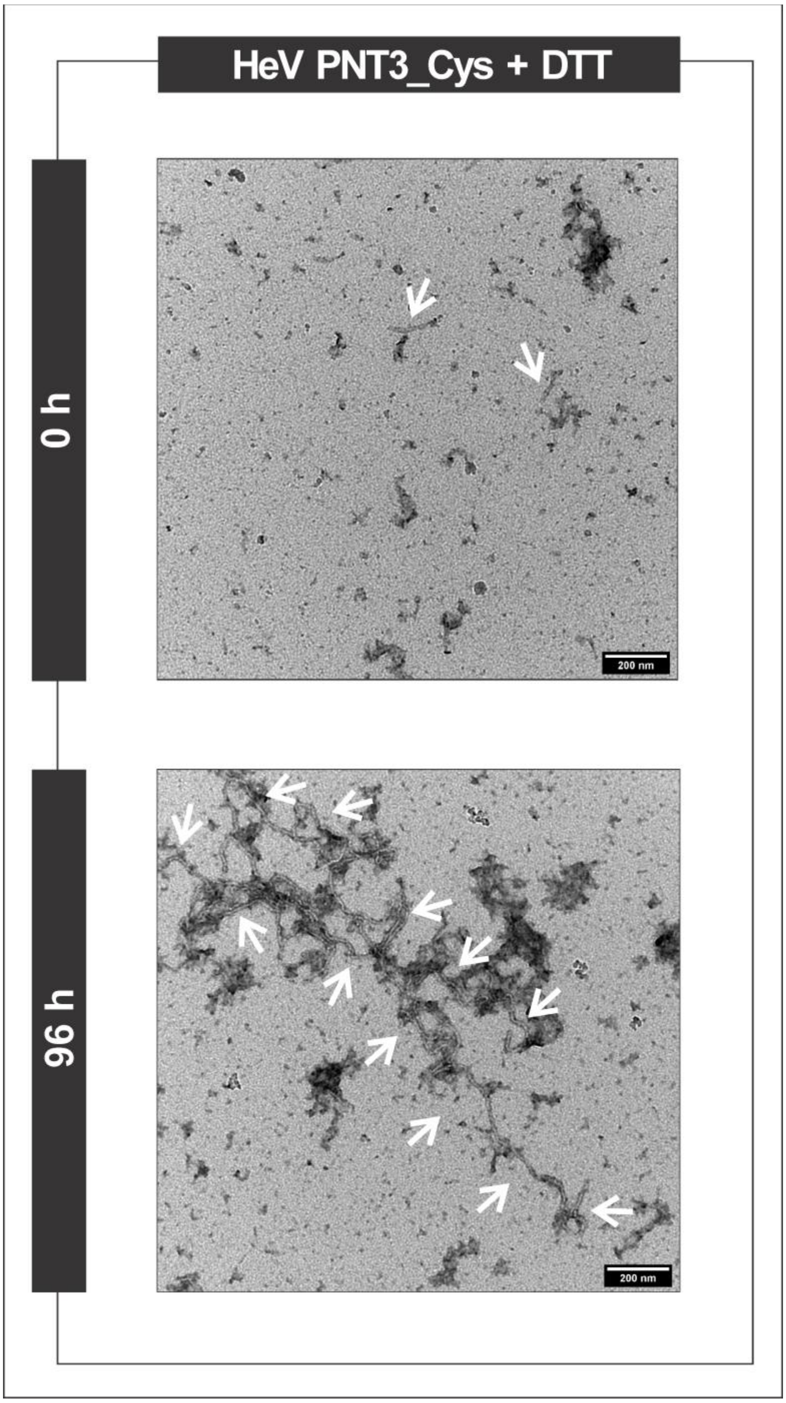


**Figure S5**. Ns-TEM analysis of HeV PNT3_Cys (200 μM) in the presence of DTT (10 mM) at time zero and after 96 h of incubation at 37 °C. Note that in all cases, samples were diluted to 40 μM prior to deposition on the grid. The white arrows indicate fibrils.
